# The genomics of linkage drag in sunflower

**DOI:** 10.1101/2022.06.07.495047

**Authors:** Kaichi Huang, Mojtaba Jahani, Jérôme Gouzy, Alexandra Legendre, Sebastien Carrere, José Miguel Lázaro-Guevara, Eric Gerardo González Segovia, Marco Todesco, Baptiste Mayjonade, Nathalie Rodde, Stéphane Cauet, Isabelle Dufau, S Evan Staton, Nicolas Pouilly, Marie-Claude Boniface, Camille Tapy, Brigitte Mangin, Alexandra Duhnen, Véronique Gautier, Charles Poncet, Cécile Donnadieu, Tali Mandel, Sariel Hübner, John M. Burke, Sonia Vautrin, Arnaud Bellec, Gregory L. Owens, Nicolas Langlade, Stéphane Muños, Loren H. Rieseberg

**Author notes:** The authors contributed equally (Kaichi Huang, Mojtaba Jahani). Joint senior authors (Nicolas Langlade, Stéphane Muños, Loren H. Rieseberg).

## Abstract

Crop wild relatives represent valuable sources of alleles for crop improvement, including adaptation to climate change and emerging diseases. However, introgressions from wild relatives might have deleterious effects on desirable traits, including yield, due to linkage drag. Here we comprehensively analyzed the genomic and phenotypic impacts of wild introgressions into cultivated sunflower to estimate the impacts of linkage drag. First, we generated new reference sequences for seven cultivated and one wild sunflower genotype, as well as improved assemblies for two additional cultivars. Next, relying on previously generated sequences from wild donor species, we identified introgressions in the cultivated reference sequences, as well as the sequence and structural variants they contain. We then used a ridge regression model to test the effects of the introgressions on phenotypic traits in the cultivated sunflower association mapping population. We found that introgression has introduced substantial sequence and structural variation into the cultivated sunflower gene pool, including > 3,000 new genes. While introgressions reduced genetic load at protein-coding sequences and positively affected traits associated with abiotic stress resistance, they mostly had negative impacts on yield and quality traits. Introgressions found at high frequency in the cultivated gene pool had larger effects than low frequency introgressions, suggesting that the former likely were targeted by artificial selection. Also, introgressions from more distantly related species were more likely to be maladaptive than those from the wild progenitor of cultivated sunflower. Thus, pre-breeding efforts should focus, as far as possible, on closely related and fully compatible wild relatives.

## Introduction

Domestication – the process that transformed wild plants into highly productive crops – is arguably the most important innovation in human history (Diamond 2002). Not only did it spark explosive population growth and the establishment of modern civilization (Diamond 1997), but it also laid the foundation for the theory of evolution (Darwin 1859) thereby unifying the life sciences (Dobzhansky 1973). While domestication and subsequent improvement have proven spectacularly successful in modifying plant architecture and enhancing yield (Evans 1993), such changes often come with a cost, including losses of genetic diversity (Tang and Knapp 2003; Khoury et al. 2022), increases in genetic load (Moyers et al. 2018), and reductions in resistance to biotic and abiotic stress (Smedegaard-Petersen and Tolstrup 1985; Mayrose et al. 2011). This is of increasing concern in the 21^st^ century, as environmentally resilient cultivars are needed to cope with a more hostile climate, while minimizing use of costly external inputs such as fertilizer, pesticides, and water.

Fortunately, diversity lost during domestication and improvement may be regained by tapping the gene pools of crop wild relatives (CWRs). The potential utility of such wild germplasm has long been recognized by plant biologists and breeders (Harlan 1975; Tanksley and McCouch 1997; McCouch et al. 2013), leading to global efforts to collect and conserve CWRs, in addition to the crops themselves (Plucknett et al. 1987). Likewise, breeding programs often include a pre-breeding component, in which wild genetic material is introduced into domesticated breeding lines (Zamir 2001; Hübner and Kantar 2021). While many such efforts have focused on enhancing disease resistance (Dempewolf et al. 2017), CWRs also have been used to increase nutritional quality, boost yield, and enhance resistance to abiotic stressors, such as drought, salt, and flooding (Gur et al. 2004; Hajjar and Hodgkin 2007; Warschefsky et al. 2014; Hübner and Kantar 2021). Economic analyses have confirmed the value of such an approach. For example, a 2013 analysis of 32 crops estimated current benefits from CWR traits to be ∼$68 billion annually, with potential future benefits of ∼$196 billion annually (PricewaterhouseCoopers 2013).

Despite the clear value of CWR traits for crop improvement, there are downsides. The introduction of wild genetic material into cultivated lines typically occurs via repeated backcrossing or introgression (Tanksley and McCouch 1997). This process is not only time-consuming, but it also can be hampered by reproductive barriers that interfere with crosses or that reduce the fitness of hybrid offspring (Moyle and Graham 2005; Tao et al. 2021). In addition, the resulting introgressions may have undesirable impacts on non-target crop traits (Chitwood-Brown et al. 2021). While this can be due to negative pleiotropic effects of the target alleles, adverse effects appear to be more frequently caused by linked alleles that are deleterious in the crop genetic background (Von fels et al. 2017; Chitwood-Brown et al. 2021), a phenomenon called linkage drag. Plant breeders typically monitor the size and location of introgressions with molecular markers and/or restrict pre-breeding efforts to fully compatible wild relatives (i.e., members of the primary gene pool; Harlan and de Wet 1971) to reduce the impact of the linkage drag (Young and Tanksley 1989; Tanksley and McCouch 1997; Frary et al. 2004). However, in large plant genomes, regions of low recombination are widespread, making it difficult to reduce the sizes of some introgressions (Rodgers-Melnick et al. 2015; Brazier and Glémin 2022; Huang et al. 2022). Also, key traits may be found outside of the primary gene pool, making it necessary to tap less compatible wild relatives (e.g., Duriez et al. 2019). The latter are classified as the secondary gene pool if they can intercross with the crop and produce at least some partially fertile hybrids (Harlan and de Wet 1971). More distantly related species that require technological interventions to produce hybrid offspring are referred to as the tertiary gene pool (Harlan and de Wet 1971).

The causes of linkage drag are assumed to be like those that contribute to species differences in natural populations. These include the genetic changes responsible for phenotypic divergence, as well as various kinds of hybrid incompatibilities (Chitwood-Brown et al. 2021; Tao et al. 2021). Introgressions with strongly negative effects are likely purged by selection during pre-breeding, so those successfully incorporated into the cultivated gene pool should be less harmful. However, as far as we are aware, a comprehensive analysis of the effects of such introgressions on cultivated phenotypes has yet to be conducted. The genomic impacts of these introgressions are even less clear. Introgression has been shown to reduce genetic load in maize (Wang et al. 2017) and sorghum (Smith et al. 2019) and to introduce gene presence/absence polymorphisms in sunflower (Owens et al. 2019), thereby increasing the size of its pan-genome (Hübner et al. 2019). However, a definitive analysis of the genomic impacts of such introgressions requires generation and analyses of multiple high-quality reference genomes.

Here we provide a comprehensive analysis of the phenotypic and genomic effects of linkage drag using sunflower as an experimental system. Crop wild relatives have been widely employed in sunflower breeding (Dempewolf et al. 2017; Seiler et al. 2017), and recent genomic studies have estimated that ca. 10% of the cultivated gene pool is derived from wild introgressions (Baute et al. 2015; Hübner et al. 2019). While most such introgressions are from wild *H. annuus*, the fully compatible progenitor of the cultivated sunflower, there are significant contributions from other species as well, making it feasible to compare the effects of introgression from the primary and secondary gene pools.

To estimate the impacts of linkage drag, we first sequenced and assembled reference genomes for seven cultivated and one wild sunflower genotype and improved the assemblies for two previously sequenced cultivars (Badouin et al. 2017). Then, using resequencing data previously generated for a diverse panel of wild donor species (Hübner et al. 2019; Todesco et al. 2020), we identified introgressions in the cultivar genomes and examined their impacts on sequence and structural variation in the cultivated sunflower gene pool. Lastly, we determined the locations of introgressions in the cultivated sunflower association mapping (SAM) population (Mandel et al. 2011) and used a ridge regression model to estimate their effects on 16 phenotypic traits, including quality traits, such as oil percentage in seeds, developmental traits such as flowering time, and yield-related traits such as head weight.

As expected, we found that introgressions increased sequence and structural polymorphism in the cultivated gene pool, reduced genetic load at protein-coding sequences, and enhanced trait values associated with abiotic stress resistance. On the other hand, introgressions typically reduced quality and yield traits. We also found that higher frequency introgressions have larger effects than low frequency introgressions, possibly indicating that the former have been targeted by artificial selection. Lastly, introgressions from the secondary gene pool had much larger negative effects than those from the primary gene pool. Thus, we encourage pre-breeding programs to focus as far as possible on the primary gene pool.

## Results

To identify SVs and introgressions in cultivated sunflowers, we constructed *de novo* genome assemblies using PacBio sequencing for seven inbred cultivated lines and one wild *H. annuus* genotype (Table 1; SI Appendix, Table S1; Dataset S1). Five of these assemblies were further scaffolded using Bionano optical mapping. We also improved the quality of previously sequenced assemblies (Badouin et al. 2017) for the HA412-HO inbred line using Illumina, 10×, and Hi-C sequencing (Table S1) and for the XRQ inbred line using the PacBio/Bionano combination described above. The nine cultivated lines represent a large part of cultivated sunflower genetic diversity present in the world’s genebanks (Terzic et al. 2020; SI Appendix, Fig. S1)

**Table 1.**
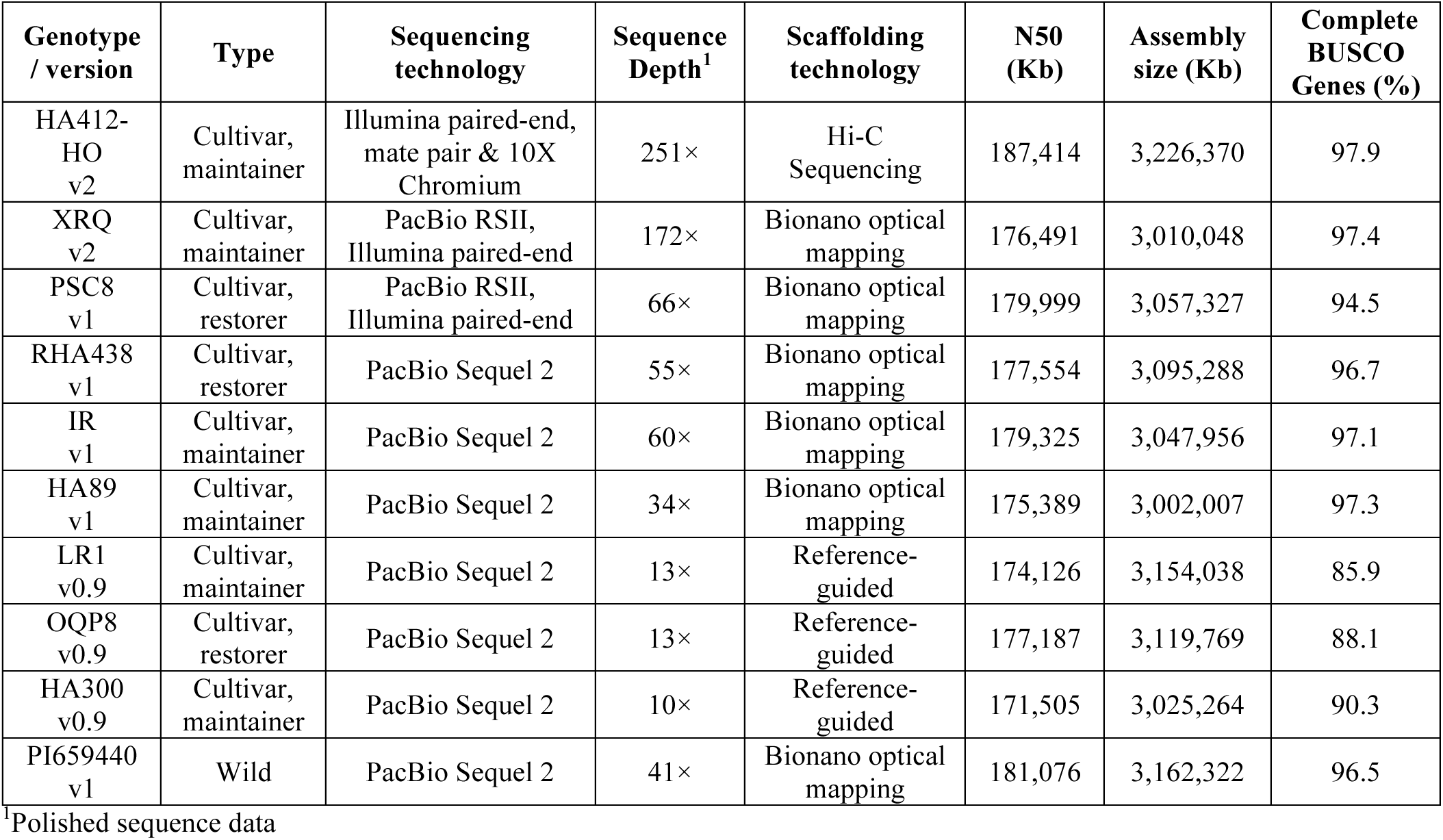
Description of new or improved reference genomes for sunflower (*H. annuus*).

| Genotype / version | Type | Sequencing technology | Sequence Depth<sup>1</sup> | Scaffolding technology | N50 (Kb) | Assembly size (Kb) | Complete BUSCO Genes (%) |
| --- | --- | --- | --- | --- | --- | --- | --- |
| HA412-HO v2 | Cultivar, maintainer | Illumina paired-end, mate pair & 10X Chromium | 251× | Hi-C Sequencing | 187,414 | 3,226,370 | 97.9 |
| XRQ v2 | Cultivar, maintainer | PacBio RSII, Illumina paired-end | 172× | Bionano optical mapping | 176,491 | 3,010,048 | 97.4 |
| PSC8 v1 | Cultivar, restorer | PacBio RSII, Illumina paired-end | 66× | Bionano optical mapping | 179,999 | 3,057,327 | 94.5 |
| RHA438 v1 | Cultivar, restorer | PacBio Sequel 2 | 55× | Bionano optical mapping | 177,554 | 3,095,288 | 96.7 |
| IR v1 | Cultivar, maintainer | PacBio Sequel 2 | 60× | Bionano optical mapping | 179,325 | 3,047,956 | 97.1 |
| HA89 v1 | Cultivar, maintainer | PacBio Sequel 2 | 34× | Bionano optical mapping | 175,389 | 3,002,007 | 97.3 |
| LR1 v0.9 | Cultivar, maintainer | PacBio Sequel 2 | 13× | Reference-guided | 174,126 | 3,154,038 | 85.9 |
| OQP8 v0.9 | Cultivar, restorer | PacBio Sequel 2 | 13× | Reference-guided | 177,187 | 3,119,769 | 88.1 |
| HA300 v0.9 | Cultivar, maintainer | PacBio Sequel 2 | 10× | Reference-guided | 171,505 | 3,025,264 | 90.3 |
| PI659440 v1 | Wild | PacBio Sequel 2 | 41× | Bionano optical mapping | 181,076 | 3,162,322 | 96.5 |

<sup>1</sup>Polished sequence data

All genomes were assembled into 17 pseudomolecules, corresponding to the 17 chromosomes in sunflower. Each of our chromosome-level genome assemblies had a total size between 3,002 and 3,226 Mb, with N50 of 172-187 Mb (Table 1; Dataset S2). The total number of genes per genome, after stringent filtering, ranged from 44,640 for XRQv2 to 63,048 genes for HA300 (Table S5). The assemblies captured 85.9-97.9% of the universally conserved single-copy benchmark (BUSCO) genes (Table 1; SI Appendix, Table S4). BUSCO percentages were positively correlated with sequence depth rather than gene number, with the lowest BUSCO scores observed for LR1 and OQP8, which were sequenced to circa 13× depth, whereas the highest BUSCO scores were seen for HA412-HOv2 and XRQv2, which were sequenced to 251× depth and 172× depth, respectively (Table 1; Dataset S1). The genomes showed high collinearity without large inter-chromosomal translocations (SI Appendix, Figs. S2-S6). Overall, our chromosome-scale genome assemblies yielded better qualitative metrics than the two previously published reference assemblies (Badouin et al. 2017).

In general, 74-83% of the genomes are composed of transposable elements (TEs), with 70-73% of these being LTR-RTs (SI Appendix, Table S6). In agreement with previous studies of the cultivated sunflower genome (Staton et al. 2012), there is a major bias in TE composition towards *Gypsy* (50-60% of total TEs) and *Copia* (13-18% of total TEs) elements, while Class II TEs (DNA transposons) were much lower in abundance relative to LTR-RTs, comprising <13% of each genome (SI Appendix, Table S6). The genomic distributions of LTR-RTs in the new assemblies are similar to those previously reported for the first reference genomes for cultivated sunflower (Badouin et al. 2017; SI Appendix, Figs. S7-15).

By mapping previously published whole-genome sequences (Hübner et al. 2019; Todesco et al. 2020) from native North American landraces (i.e., early domesticates) and five wild possible donor species to each genome assembly, we determined the ancestry of each cultivated line and estimated the locations and likely parentage of introgressions. Only a small portion (2-8%) of each genome was admixed (Fig. 1; SI Appendix, Fig. S16; Dataset S3), which is similar to previous estimates for the XRQ and HA412-HO genomes (Badouin et al. 2017). All cultivated genomes possessed more introgressions from the primary gene pool (primary introgressions) than those from the secondary gene pool (secondary introgressions).

**Fig. 1.**
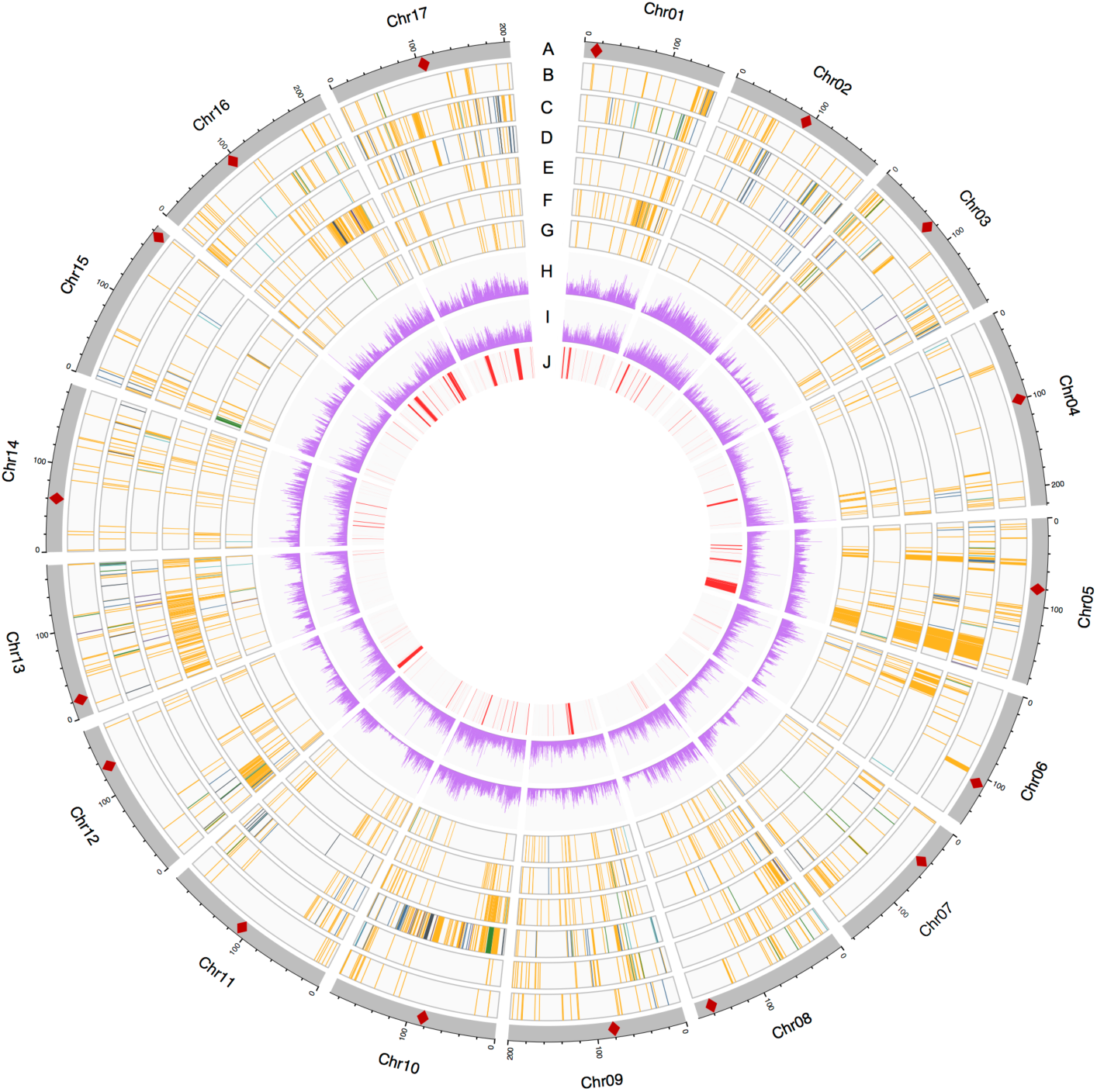
Introgressions and genetic variants of the high-contiguity cultivated sunflower genome assemblies. **A.** Chromosomes of the HA412-HOv2 reference. Diamonds mark approximate positions of centromeres. **B-G.** Introgressions in HA412-HO, XRQ, PSC8, RHA438, IR, and HA89 projected to the Ha412-HOv2 reference. Colored bars represent introgressions from different wild donors: orange: *Helianthus annuus*, green: *H. argophyllus*, light blue: *H. petiolaris* subsp. *petiolaris*, deep blue: *H. petiolaris* subsp. *fallax*, purple: *H. niveus* and dark grey: *H. debilis*. **H-I.** Density of SNPs (**H**) and small InDels (**I**) (number/500 kb; 0-10000 for SNPs and 0-2000 for small InDels). **J.** Inversions identified in genome assemblies. Regions of introgression less than 1 Mb were thickened to 1Mb for visualization.

Sunflower is a hybrid crop, and crop wild relatives were used to develop cytoplasmic male sterile “female” lines and branching, fertility restoring “male” lines for hybrid production. The male restorer lines PSC8, OQP8, and RHA438 generally had more introgressions than the female maintainer lines (HA412, XRQ, IR, HA89, LR1, and HA300). Consistent with breeding records and previous findings (Gentzbittel et al. 1999; Baute et al. 2015; Vear 2016; Hübner et al. 2019), the restorer lines had substantial introgression from wild *H. annuus* on chr10, which underlies apical branching, as well as an introgression near the distal end of chr13, where the restorer of fertility locus (*Rf1*) of the common PET1 male sterile cytoplasm is located (Fig. 1). However, while the restorer allele in PSC8 and OQP8 was derived from *H. petiolaris* as expected (Leclercq 1969), an introgression from wild *H. annuus* was found in RHA438 at the region, suggesting possible different origins of fertility restoration in cultivated sunflower. The majority (∼68%) of the primary introgressions were unique to one genotype and only a small proportion (<0.1%) were shared across all nine genomes. Almost all secondary introgressions were unique to one genotype.

We identified single nucleotide polymorphisms (SNPs) and small (<50bp) insertions/deletions (InDels), as well as different types of structural variants (SVs) including large (> 50 bp) InDels, copy number variants (CNVs), inversions, and translocations through the alignment of the high-contiguity cultivar genome assemblies (HA412-HOv2, XRQv2, PSC8, RHA438, IR, HA89). In total, we identified 12,036,913 SNPs and 3,005,855 small InDels across 17 chromosomes using the HA412-HOv2 genome as the reference (Fig. 1). We also detected 70,612-84,709 large InDels, 32,668-47,706 CNVs, 4,776-7,738 translocations, and 261-301 inversions (>1kb) between each genome and the HA412-HOv2 reference (Fig. 1; Dataset S4). After merging, 532 polymorphic inversions with a total size of 200 Mb were identified across the cultivars, including a 21-Mb region (156-177Mb) on chr5 that corresponded to the largest section of a cluster of inversions previously identified in wild *Helianthus annuus* (Todesco et al. 2020; Fig. 1J).

### Introgression Introduced Substantial Sequence and Structural Variation into the Cultivated Sunflower Gene Pool

We compared densities of SNPs and small InDels between regions with introgression in one to five genomes (polymorphic introgressed regions) and those without introgression in any of the six highly contiguous cultivar genomes (non-introgressed). We calculated densities of SNPs and small InDels in non-overlapping windows of 500kb using the HA412-HOv2 genome as the reference and compared between polymorphic introgressed regions and non-introgressed regions. Overall, regions polymorphic for primary or secondary introgressions had more SNPs and small InDels than non-introgressed regions (Fig. 2A,B). Secondary introgressions had more SNPs and small InDels than primary introgressions, although the differences were not significant. Analyses of 287 individuals comprising the cultivated SAM population (see below) revealed that introgressed regions also possessed significantly higher numbers of SNPs compared to non-introgressed regions, and secondary introgressions displayed significantly more SNPs than primary introgressions (SI Appendix, Fig. S17).

**Fig. 2.**
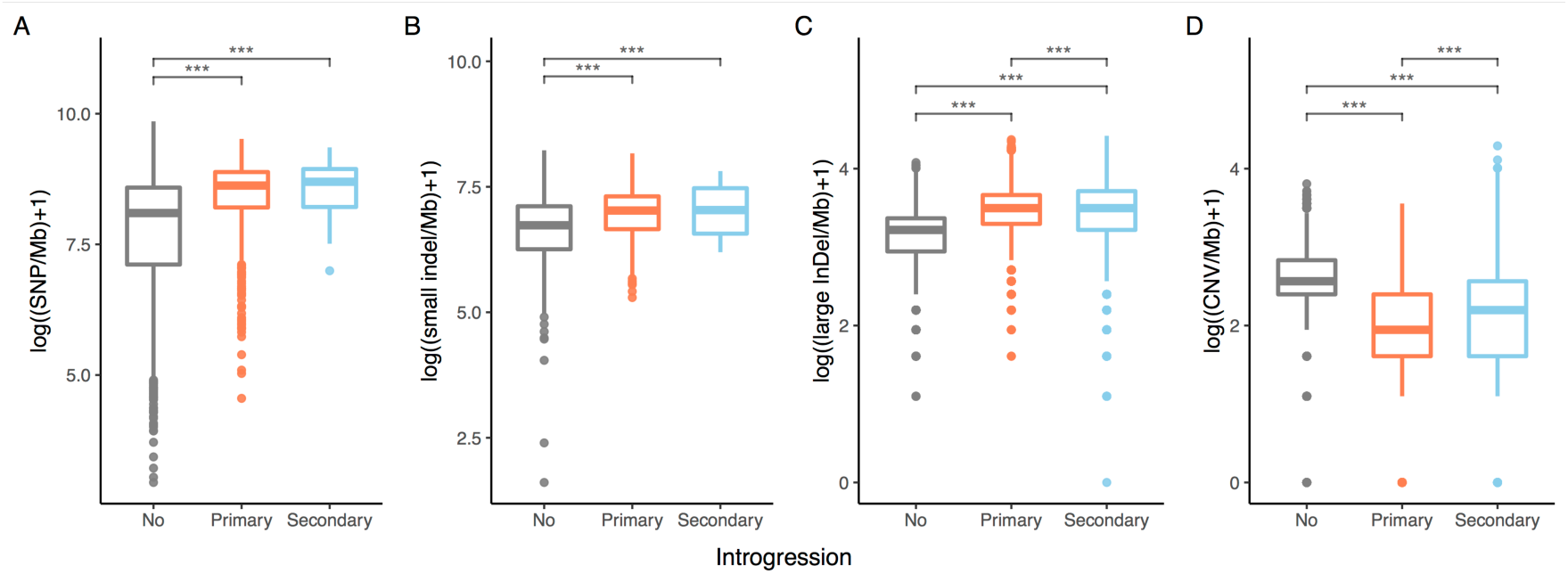
Densities of **A.** SNPs, **B.** Small InDels (<50bp), **C.** Large InDels (>50bp) and **D.** CNVs in regions without introgression, regions with introgressions from the primary gene pool (primary introgressions) and regions from the secondary gene pool (secondary introgression). The densities of SNPs and small InDels were calculated in non-overlapping windows of 500kb using the HA412-HOv2 genome as the reference. Densities of large InDels and CNVs were calculated in 10,000 samplings of 500kb windows in each type of region between each genome and the HA412-HOv2 reference. Asterisks denote significance in independent t-tests: \*\*\**P*<0.001.

Wild introgressions also introduced large (>50bp) insertions and deletions (large InDels) into the cultivated sunflower gene pool. In each pair of genome comparisons with the HA412-HOv2 reference, both primary and secondary introgressions had significantly higher numbers of large InDels compared to regions without introgression (Fig. 2C). Conversely, introgressions had significantly fewer CNVs than non-introgressed regions (Fig. 2D). We suspect that this is due to the reduced strength of purifying selection on TE copy number in the cultivated gene pool.

Across the six high-contiguity genomes, chromosomal inversions had an overlap of 58 Mb with primary introgressions and 5.7 Mb with secondary introgressions, which is significantly higher than a random distribution in both cases (primary introgressions: *P*<0.001, secondary introgressions: *P*=0.0269). In each pair of genome comparisons with the HA412-HOv2 reference, the number of inversions introduced from the primary gene pool varied from 0.24 to 0.43 per Mb, which is significantly (*P*<0.01) higher than that in non-introgressed regions (0.07-0.08/Mb). More inversions were introduced from the secondary than from the primary gene pool in each genome, except in HA89 where no inversions were found in secondary introgressions (SI Appendix, Fig. S18).

### Introgression Reduced Genetic Load

We estimated the effect of introgression on genetic load by calculating the ratio of the number of alternative stop codons (*P*_nonsense_) and the number of nonsynonymous mutations (*P*_nonsyn_) in 500-kb sliding windows (Renaut and Rieseberg 2015). The statistic was negatively correlated with recombination rate (SI Appendix, Fig. S19), in accord with previous understanding of the role of recombination in eliminating deleterious mutations (Huang et al. 2022). *P*_nonsense_/*P*_nonsyn_ of polymorphic primary introgressions was lower in null recombination rate regions than that of non-introgressed regions and comparable to non-introgressed regions in regions of reduced and high recombination rate (SI Appendix, Fig. S19). Secondary introgressions displayed a trend towards reduced load (i.e., lower *P*_nonsense_/*P*_nonsyn_ ratios) compared to non-introgressed regions, but the sample size was too small to draw conclusions. Analyses of 287 individuals in the cultivated SAM population (see below) provided clearer results. While *P*_nonsense_/*P*_nonsyn_ was also negatively correlated with recombination rate in this dataset (SI Appendix, Fig. S20), primary introgressions displayed significantly lower *P*_nonsense_/*P*_nonsyn_ than non-introgressed regions in all recombination rate categories, and secondary introgressions had significantly lower *P*_nonsense_/*P*_nonsyn_ than non-introgressed regions in regions of null and reduced recombination rate (Fig. 3).

**Fig. 3.**
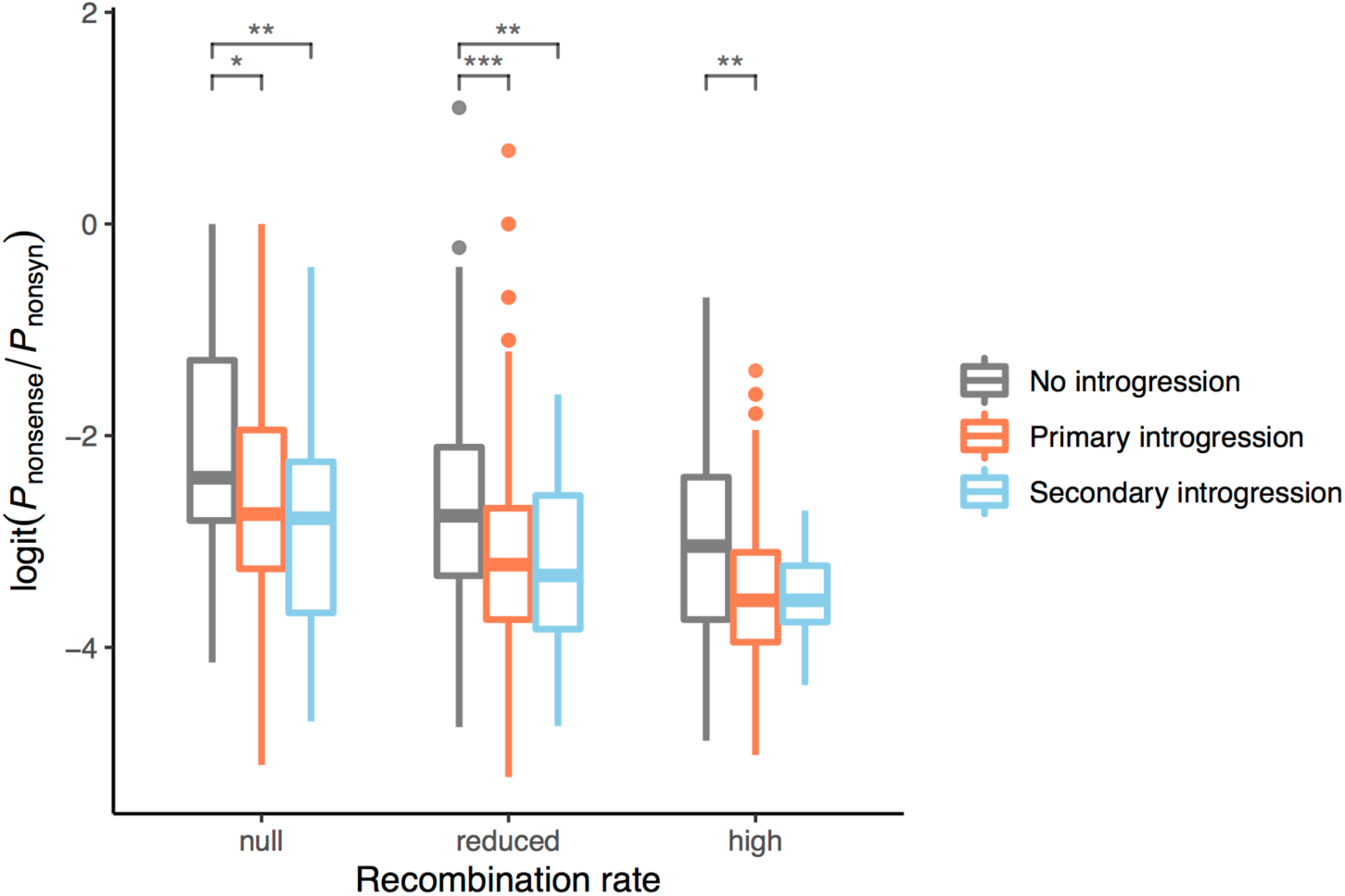
Ratio of alternative stop codons and nonsynonymous mutations (*P*_nonsense_/*P*_nonsyn_) in regions without introgression, regions with introgressions from the primary gene pool (primary introgressions) and regions from the secondary gene pool (secondary introgressions) in the cultivated sunflower association mapping population. *P*_nonsense_/*P*_nonsyn_ was calculated in non-overlapping windows of 500kb. Windows of each recombination rate category (high: > 2 cM/Mb, reduced: 0.01-2 cM/Mb, null: <0.01 cM/Mb) were compared separately. Asterisks denote significance in independent t-test: *0.05>*P*>0.01, **0.01>*P*>0.001, \*\*\**P*<0.001.

### Introgressions Introduced Gene Presence/absence Polymorphisms

A total of 77,334 genes were obtained across the 10 genome assemblies, among which 75,791 were present in the 9 genomes of cultivars. Altogether, 31,099 genes in the pan-genome displayed PAV between genomes. After filtering based on synteny, we retained 75,369 genes with coordinate information for homologs, 29,948 of which showed PAV.

We found that introgressions introduced significantly more gene PAVs than non-introgressed regions, but gene PAVs from primary and secondary introgressions did not differ significantly, except in one pair (Fig. 4). The total number of genes introduced by primary introgressions ranged from 889 for HA300 to 4,323 for RHA438, respectively, whereas between 26 (HA89) and 1,800 (OQP8) genes were introduced by secondary introgressions (SI Appendix, Fig. S21). On average, 12% of the PAVs result from primary introgressions and 5% from secondary introgressions. Across the nine cultivar genomes, a total of 3,187 genes were introduced by introgression from crop wild relatives. Unsurprisingly, the number of new genes introduced by introgression is closely correlated with total amount of introgression detected in a genome, so we see more new genes resulting from introgression in the restorer lines (PSC8, RHA438 and OQP8) than from maintainer lines (SI Appendix, Fig. S21).

**Fig. 4.**
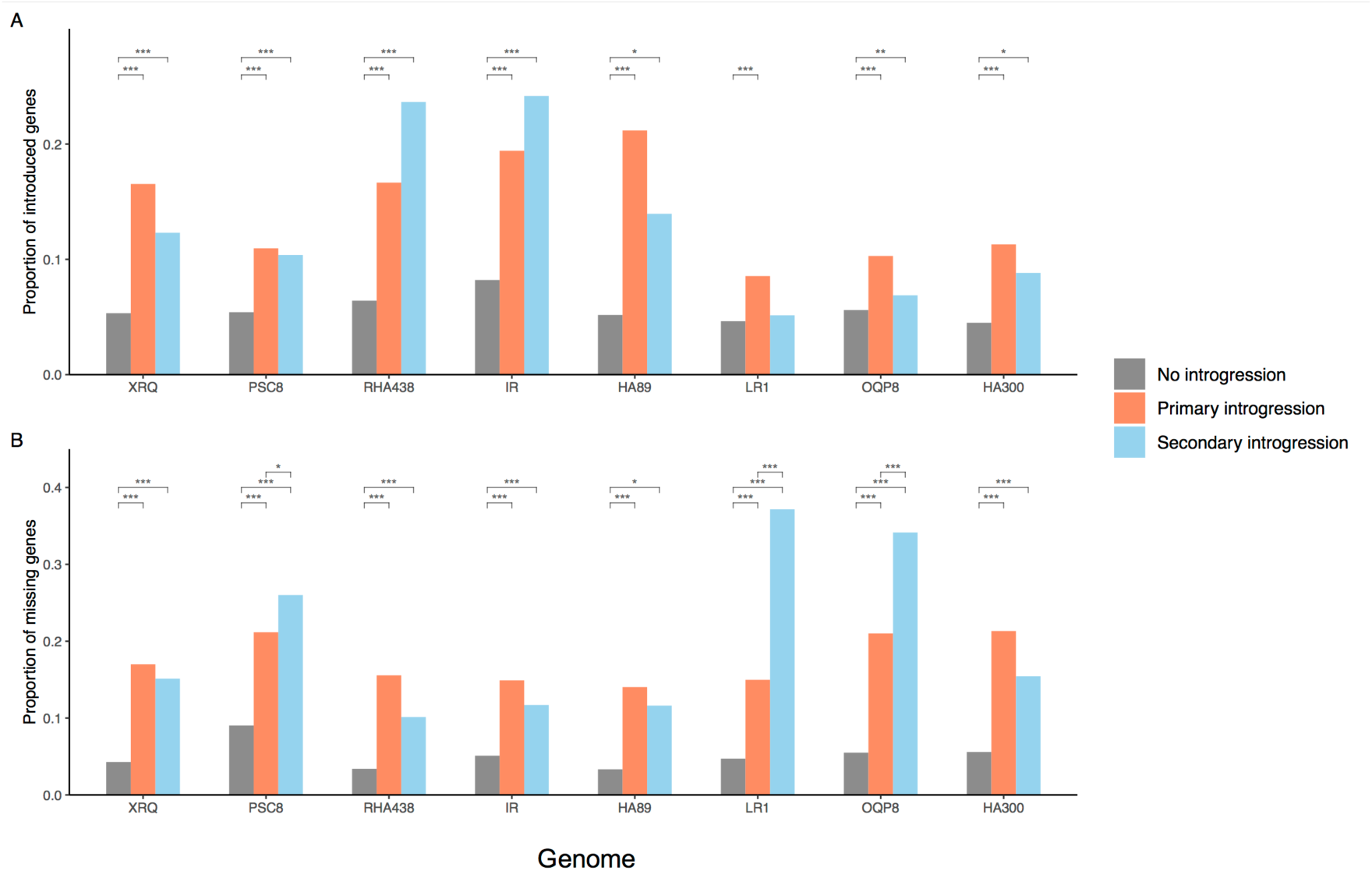
Proportions of **A.** introduced genes and **B.** missing genes in introgressed and non-introgressed regions in each cultivar genome compared to the HA412-HOv2 reference. Asterisks denote significance in independent t-test: *0.05>*P*>0.01, **0.01>*P*>0.001, \*\*\**P*<0.001.

In addition to new genes, introgressions often lack genes that are present in syntenic non-introgressed regions (Fig. 4B). Primary introgressions introduced 383 (HA300) to 1,577 (RHA438) missing genes, whereas between 22 (HA89) and 2095 (OQP8) gene absences were caused by secondary introgressions (SI Appendix, Fig. S21). About 17-32% of the gene absences in primary introgressions had a homolog present in the wild *H. annuus* (PI659440) genome, indicating that many of such missing genes represent gene PAVs in the wild donor species.

### Introgressions in the Cultivated Sunflower Association Mapping (SAM) Population

We generated a SNP dataset using previously published sequence data for 287 individuals in the SAM population (Mandel et al. 2011; Hübner et al. 2019), as well as the aforementioned whole-genome sequences from native North American landraces and five possible wild donor species. Then, we determined the locations and parentage of introgressions in each of the 287 cultivated genotypes. We found that all samples contained putative introgressions and that all chromosomes appeared to have experienced introgression in at least one of the SAM samples (Fig. 5). The amount of introgression in each sample varied from 0.4% to 11% with a number of samples having large blocks of introgression (Dataset S5). On average, each sample had ca. 3% of the genome covered with introgressions from the primary gene pool and 0.1% derived from the secondary gene pool, which is similar to the estimates from the genome assemblies, but lower than previously estimated for the SAM population using a different method (Hübner et al. 2019). Restorer lines had more introgression than maintainer lines on average (3.8% vs. 2.9%). Maintainer and restorer lines showed distinct patterns of introgression on the first half of chr8, a substantial portion of chr10, part of chr12, as well as the end of chr13, broadly consistent with previously identified regions of high divergence between these groups (Baute et al. 2015; Hübner et al. 2019; Owens et al. 2019). Small regions of introgression from the secondary gene pool were identified at the end of chr13 in most of the restorer lines, but not in maintainers. These regions roughly correspond to the introgression from *H. petiolaris* in the PSC8 genome, corroborating previous findings of the *Rf1* restorer allele at this position (Gentzbittel et al. 1999; Baute et al. 2015).

**Fig. 5.**
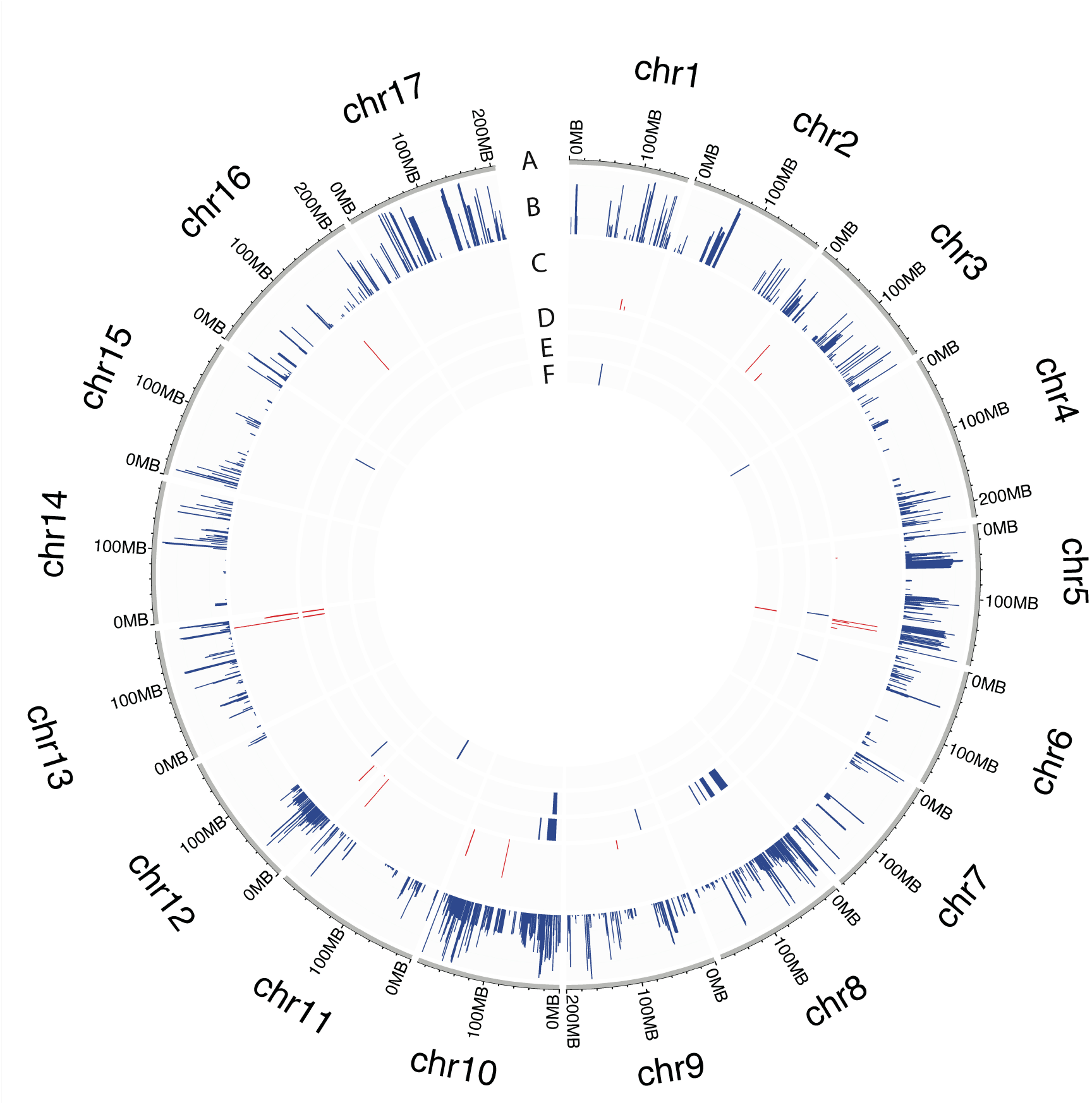
Frequency of introgression variants in the SAM population and associated introgressions with traits in GWA analysis. **A.** Chromosomes of the HA412-HOv2 reference. **B.** Frequency of introgression variants from the primary germplasm. **C.** Frequency of introgression variants from secondary germplasm. **D.** Introgressed genomic intervals associated with developmental traits (number of branches, head weight, head diameter, stem weight, leaf weight, and plant biomass). **E.** Introgressed genomic intervals associated with quality traits (seed size and oil percentage). **F.** Introgressed genomic intervals associated with flower pigmentation (anthocyanins in disk florets, anthocyanins in stigmas). Blue and red representing introgressions from primary and secondary germplasm, respectively.

Using these datasets, we evaluated the presence or absence of introgressions in 1kb non-overlapping windows across the genome. We took this approach to account for the fact that most introgressions are fragmented by recombination as they are incorporated in the cultivated sunflower gene pool and to permit genome wide association studies (GWAS) and various population genomic analyses. A total of 505,038 and 5,243 introgression variants were detected at a ≥ 3% minor allele frequency cut off for primary (wild *H. annuus*) and secondary germplasm donors, respectively (Fig. 5).

We then performed GWAS of the introgression variants for 16 traits that were previously phenotyped (Mandel et al. 2013; Nambeesan et al. 2015; Lee et al. 2022) in common gardens at three locations (Watkinsville, GA, Ames, IA, and Vancouver, BC) using a model that corrects the population structure and familial relatedness. Our results revealed that introgressions have a significant effect on the phenotypic variation in the SAM population (SI Appendix, Fig. S24). After merging GWA outliers in the range of 10 Mb, introgression intervals were found to underlie 27 quantitative trait loci (QTLs) for 12 phenotypic traits (Table S7; Fig. 5). Of these, 23 (85.18%) were introgressed from primary germplasm (wild *H*. *annuus*), while 4 (14.81%) were introgressed from secondary germplasm. The introgressed QTLs reduced head diameter and head weight, but increased plant biomass, number of branches, anthocyanins in disk florets, number of days to flowering, dry leaf weight, oil percentage, seed size, dry stem weight, and anthocyanins in stigmas. For stem diameter, introgressed QTLs with negative and positive effects were found. The 27 QTLs were not fully independent. A primary introgression near the beginning of chr10 that introduced branching into restorer lines, also effects oil content, seed size, head diameter, and head weight.

However, GWAS does not consider the effects of introgression variants that fall below a stringent significance threshold. Therefore, we employed the following ridge regression model to estimate phenotypic effects across all introgression variants:

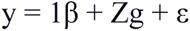

Where y is a vector of the phenotypic trait; Z is an incidence matrix containing the allelic states of the markers (Z = {-1, 1, 0}); -1 and 1 represent homozygous non-introgressed and introgressed genotypes at a locus, respectively and 0 represents the heterozygous state; β is a vector of fixed effects; g is the vector of marker effects; and ε is a vector of residuals.

To assess whether introgressions overall have a significant impact on the 16 phenotypic traits, we compared the average value of introgression marker effects to a null distribution. Our results indicated that introgressions overall have negative effects on traits associated with yield, including head diameter, head weight, leaf area, leaf weight, seed size, seed weight, stem diameter, and stem weight (Fig. 6). This pattern was seen for introgressions from both the primary (wild *H. annuus*) and secondary gene pool. In contrast, biomass, branching, and specific leaf area (SLA) showed an increase in the trait value for introgressions from both gene pools. Branching was introgressed into restorer lines to prolong the flowering period for hybrid production and increased SLA is thought to be associated with drought tolerance (Wellstein et al. 2017), so both changes can be viewed as potentially desirable. We also observed gene pool-specific effects for stigma and disk floret anthocyanins and oil percentage; primary introgressions increase anthocyanin content and reduce oil percentage, whereas introgressions from secondary germplasm do not cause significant change (Fig. 6). Lastly, a comparison of effect sizes of introgression variants from the primary versus secondary gene pool indicate that the latter have much larger effects on average (SI Appendix, Fig. S25).

**Fig. 6.**
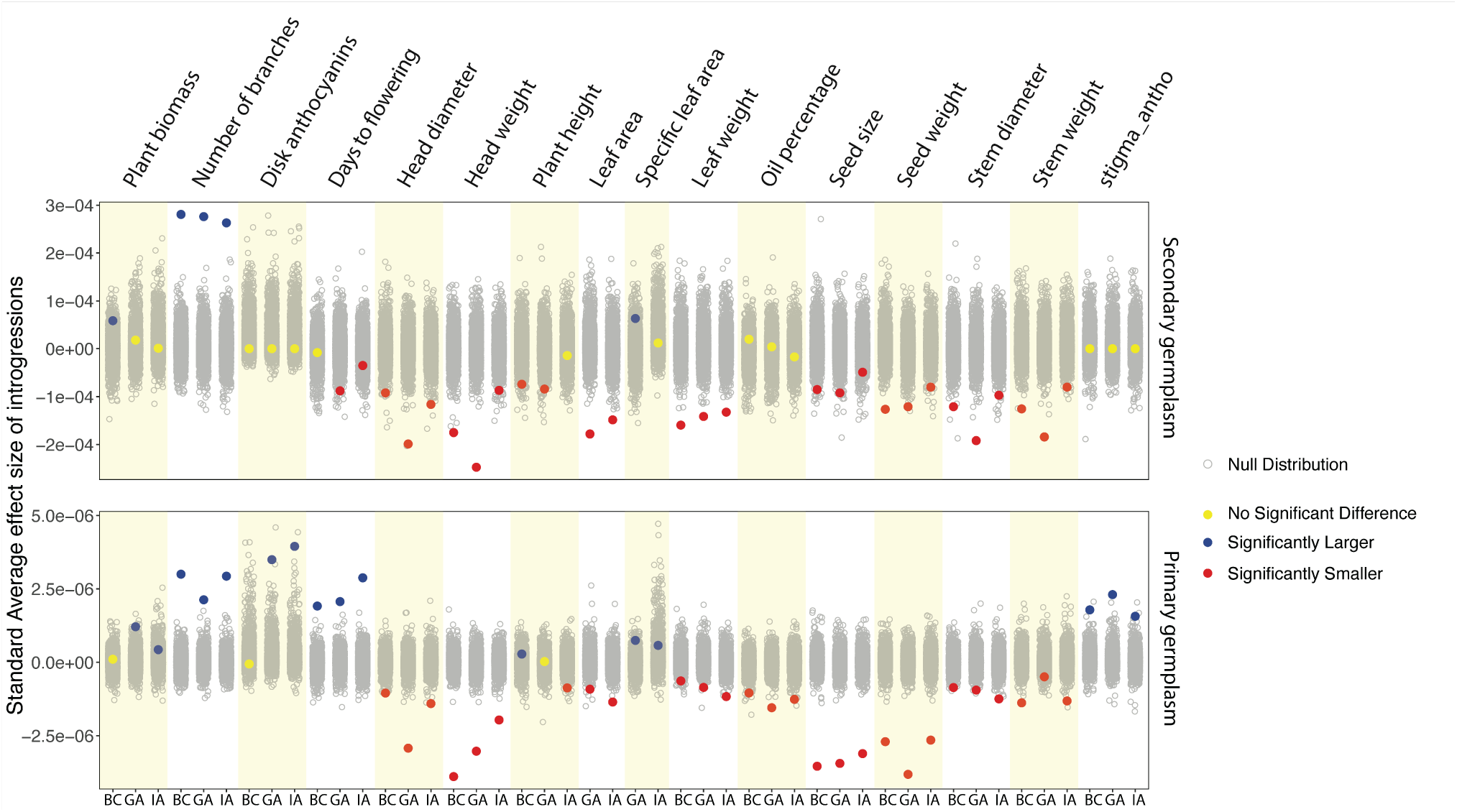
Standardized average effect sizes of introgression variants (calculated from Z-score normalized trait values) in the SAM population. Gray dots show the null distribution of effect sizes. Red, blue and yellow represent the decreasing, increasing, or neutral effects of introgressions on phenotypic traits at P-value < 0.05. BC: British Columbia, GA: Georgia, IA: Iowa.

Next, we asked whether the frequency of introgression variants was correlated with their effect size. Higher frequency introgressions are more likely to have been targets of artificial selection, so we were especially interested in the potential for linkage drag associated with such introgressions. We found a significant correlation (*P*<0.05) between the frequency and the effect size of introgression variants from both the primary and secondary gene pools across all traits and common gardens (Fig. 7; SI Appendix, Fig. S26). In general, higher frequency introgressions have larger phenotypic effects than lower frequency introgressions. Changes in beta coefficients were mostly consistent between donor gene pools: biomass, branching, SLA, oil percentage, and stigmas anthocyanins had positive beta values for introgressions from both the primary and secondary gene pool, whereas negative beta values were observed for the other traits.

**Fig. 7.**
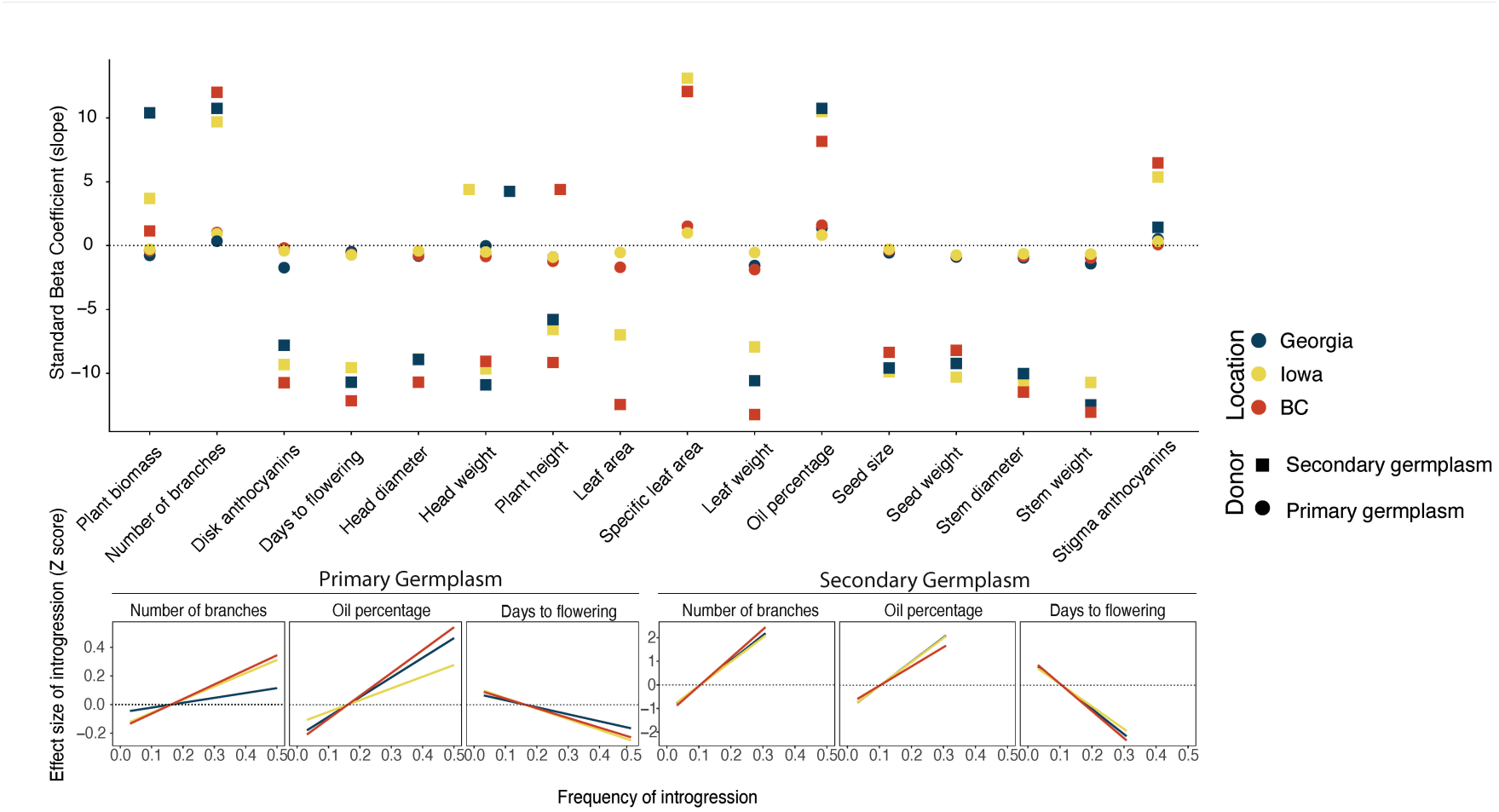
Results from linear regression model where X = introgression frequency in SAM population and Y = introgression effect on phenotype trait. **A.** The standard beta coefficient of all traits in three common garden experiments. **B.** A fitted linear regression line for branching, oil percentage, and days to flower for introgressions from primary and secondary germplasm.

## Discussion

### New Genomic Resources for Sunflower

For the past two decades, the plant biology community has made substantial investments into the generation of genomic tools and resources for crops and their wild relatives, especially high-quality reference sequences (Thudi et al. 2021). These investments are now bearing dividends, ranging from exciting new discoveries about plant domestication (Purugganan 2021) to the genetic dissection of key ecological and agronomic traits (Kuroha et al. 2018; Temme et al. 2020) to increases in the speed and precision of plant breeding (Jannink et al. 2010). Despite these successes, the goalposts have moved. Plant genomes have been shown to vary remarkably in their content and structure, even within species (Lei et al. 2021; Bayer et al. 2020), and these differences often underlie variation in phenotypic traits (Gage et al. 2019). Thus, tens or even hundreds of reference quality genomes are needed to fully understand the genomic basis of phenotypic variation (Gage et al. 2019; Della Coletta et al. 2021). Here we report progress toward this goal by providing eight new chromosome-level genomes for sunflower along with significant improvements of two previously published sunflower genomes (Badouin et al. 2017). These ten publicly available genomes, which encompass much of the genetic space in the cultivated sunflower gene pool (SI Appendix, Fig. S1), represent a valuable resource for sunflower research and breeding.

While the genomes were sequenced and assembled using different sequencing technologies and depths, we were able to obtain chromosome level assemblies for all genotypes, even with sequencing depth as low as 10× when using PacBio HiFi reads and reference-guided assembly (for HA300; Table 1). We did see a trade-off between lower sequence coverage and BUSCO scores, suggesting that the quality of gene annotation suffers at lower sequencing depths. However, excellent BUSCO scores were obtained with sequence depth in the 30x range with HiFi reads, which may represent an optimal balance between sequencing cost and genome quality.

The cultivated genomes range from 3.02 to 3.23 Gb in size, with the wild genome at 3.16 Gb falling in the middle. Thus, domestication in sunflower does not appear to been accompanied by a change in genome size. On the other hand, the 10 genomes are ca. 15% smaller than previous genome size estimates for *H. annuus* (which included HA89, one of the genomes sequenced here) based on Feulgen staining (Sims and Price 1995) and flow cytometry (Baack et al. 2005). Given that the two different scaffolding approaches (Bionano optical mapping and Hi-C sequencing) employed in the present study resulted in similar genome size estimates, we suspect that previous work over-estimated the size of the sunflower genome.

Synteny comparisons of the six high-contiguity genomes failed to reveal large-scale chromosomal rearrangements between the genomes, except for one 21 Mb inversion. However, we did find millions of small indels, thousands of deletions and insertions, and hundreds of inversions. We also detected numerous differences in gene content, with approximately 40% of the 77,334 genes in the sunflower pan-genome varying between genomes. This is higher than the 27% previously reported based on re-sequencing data from the SAM population (Hübner et al. 2019), possibly because the present study is based on comparisons of fully assembled reference genomes. Estimates of the proportion of genes displaying presence absence polymorphisms in other plant species range from 15-66% (Bayer et al. 2020; Hufford et al. 2021), so the level of polymorphism in sunflower is not unusual. Like other plant species, gene presence-absence polymorphisms have been shown to play an important functional role in sunflower. For example, Todesco et al. (2020) showed that a PAV for *HaFT1* was responsible for a 77-day shift in flowering time between two ecotypes of the silverleaf sunflower. More recently, Lee et al. (2022) found that the complementation of PAVs in sunflower hybrids was the primary cause of heterosis.

### Genomic Consequences of Introgression

Analyses of the ten genomes provide insights regarding the sources of variation among them. Consistent with previous reports, about three quarters of the sunflower genome is made up of LTR transposons and other TEs, and many of the differences between genomes result from variability in the accumulation, movement, and elimination of TEs (Badouin et al. 2017). Also, sunflower is the product of a whole genome duplication event approximately 29 Mya (Barker et al. 2008, 2016; Badouin et al. 2017), and the differential retention of duplicated sequences likely contributes to genomic diversity as well.

Introgression from wild relatives represents another potential source of variation (Hübner et al. 2019; Owens et al. 2019). By examining the location and parentage of introgressions in the cultivated genomes, we were able to show that introgressed regions have greater diversity than non-introgressed regions as measured in terms of SNPs, small indels, deletions, insertions, inversions, and gene PAVs. The impact of the introgressions was most pronounced for the latter, with introgressions accounting for about 17% of PAVs. This is qualitatively similar to wheat, where differences in the gene content of introgressions from divergent donors appears to cause reduced performance (Hao et al. 2020). Introgressions also reduced genetic load at protein coding genes and variation in CNVs, possibly because of relaxed purifying selection in the cultivated gene pool. CNVs in sunflower are mostly caused by variation in TE copy number, which may explain why introgression affects them differently than gene PAVs.

A previous study of the SAM population showed that the absence allele at PAVs often has deleterious impacts on yield-associated traits (Lee et al. 2022), and we speculate that they may be the primary genetic cause of linkage drag. The genetic architecture of linkage drag has implications for mitigation strategies. If the maladaptive allele is commonly the absence variant of a PAV, then it could be complemented in hybrids containing the domesticated allele, whereas an allele that was maladaptive for other reasons (e.g., additive effect polygenes) is unlikely to be rescued in hybrids. Unfortunately, we were unable to directly test this hypothesis in the present study because the SAM population is comprised mainly of inbred lines.

### Phenotypic Consequences of Introgression

Introgressions from the primary gene pool (i.e., wild *H. annuus*) had a significant impact on all 16 traits phenotyped in the SAM population, whereas those from the secondary gene pool affected 13 of the 16 traits (Fig. 8). This is unsurprising since introgressions from wild *H. annuus* are much more frequent in the SAM population than those from the secondary gene pool. On the other hand, the effect sizes of secondary introgressions are much larger on average than those from wild *H. annuus* (SI Appendix, Fig. S25).

Examination of the direction of effects of the introgressions indicates that most reduce desirable agronomic trait values, especially traits that correlate closely with yield, including head diameter, head weight, seed size, and seed weight, though there are exceptions. For example, introgressions typically increase SLA, which is frequently associated with greater drought tolerance (Wellstein et al. 2017). This makes sense given that sunflower wild relatives are more drought tolerant than cultivars (Baack et al. 2008; Seiler et al. 2017). In addition, introgressions show an increase in biomass, but this appears to be a by-product of increased branching, which has been introduced into restorer lines to prolong flowering and thus pollen shed. Lastly, while introgressions may negatively affect traits on average, there can be individual introgressions with effects in the opposite direction. For example, an introgression on chr10 from wild *H. annuus* that is associated with increased branching also results in increased oil content and seed size (Table S7). Overall, however, introgressions from wild *H. annuus* negatively affected the latter two traits.

An unexpected result was that higher frequency introgressions had larger effects on traits (both positive and negative). We speculate that such high frequency introgressions have been directly targeted by artificial selection. In some instances, the trait we phenotyped was likely the target of selection (e.g., branching and oil content), whereas maladaptive trait values are most likely the product of linkage drag for traits such as disease resistance that were not phenotyped in the present study.

## Conclusions

In summary, by utilizing a combination of high-quality reference genomes and genotypic and phenotypic analyses of the SAM population, we provide a comprehensive assessment of the impact of linkage drag on the cultivated sunflower genome and on plant performance. We show that despite the numerous benefits deriving from tapping crop wild relatives, such as the introduction of desirable traits and genetic and phenotypic variation (Warschefsky et al. 2014; Dempewolf et al. 2017), there can be downsides, including reductions in yield-related traits. We speculate that this is largely due to the introduction of variation in gene content; cultivars containing introgressions not only have new genes, but they also are missing genes that would otherwise be present, which can have deleterious consequences (Lee et al. 2022).

So, what strategies can be employed to mitigate the effects of linkage drag? Marker-assisted selection is widely employed to reduce the sizes of introgressed regions (Young and Tanksley 1989; Hao et al. 2020), although this can be challenging in genomic regions of low recombination, such as near the branching locus on chr10. Genome editing and other biotechnology approaches have the potential to introduce favorable alleles without linkage drag (Kawall 2019), although we recognize that the application of such approaches are currently limited by regulatory and socio-political factors (Friedrichs et al. 2019). If the genetic factors underlying linkage drag are mostly recessive, such as would be the case for missing genes, then hybrid production offers an effective strategy for ameliorating linkage drag. Lastly, our results indicate that introgressions from distantly related species are much more problematic than those from the fully compatible wild progenitor of cultivated sunflower. Thus, linkage drag could be ameliorated by restricting pre-breeding efforts to closely related and fully compatible wild relatives. While certain desirable traits might not be expressed in close relatives, many of the underlying alleles may exist in the primary gene pool, albeit at a lower frequency. If so, there is a growing potential for the use of bioinformatics approaches to identify compatible genebank germplasm containing the allele(s) of interest (Guerra et al. 2022). Furthermore, natural introgression from the secondary gene pool into the primary gene pool may provide a source of alleles that have already been purged of deleterious incompatibilities and show reduced linkage drag.

## Materials and Methods

For full materials and methods, see SI Appendix, Supplementary Information Text.

### Diversity Analyses

To show the relationships of the nine sequenced inbred lines to cultivated sunflower genetic diversity, we positioned them in genetic space using principal components analysis (SI Appendix, Fig. S1) based on unpublished genotypic data comprising 16,048 SNP markers genotyped on 2,850 cultivated lines.

### Nucleic Acid Extractions, Library Preparations, and Sequencing

For DNA sequencing, high molecular weight DNA was extracted from young leaves using several different protocols, including a modified CTAB protocol (Todesco et al. 2020) for HA412-HO, magnetic bead extraction (Mayjonade et al. 2016) for the remaining cultivated genotypes, and the QIAGEN Genomic-tip 100g procedure for PI659440.

For the HA412-HOv2 genome (which is an updated version of the HA412-HO genome, Badouin et al. 2017), paired-end and mate-pair libraries were generated and sequenced using Illumina sequencing technology to a total depth of 214× (Dataset S1). In addition, 10× Genomics Chromium libraries were prepared and sequenced using Illumina to 37× depth (Dataset S1).

For XRQv2 (which is an updated version of the XRQ genome; Badouin et al. 2017) and the newly sequenced genotypes, library preparation and sequencing employed Pacific Biosystems (PacBio) technology (Dataset S1). RSII system raw reads were generated for XRQv2 and PSC8, Sequel II system raw/CLR plus HiFi reads for IR and RHA438, and Sequel II HiFi reads for PI659440, HA89, LR1, OQP9 and HA300.

We sequenced full-length cDNA using PacBio SMRT sequencing technology (IsoSeq) for the IR, RHA438, PI659440, and HA89 lines. In brief, leaf, bud and stem tissues were collected for each accession, flash frozen in liquid nitrogen. RNA was subsequently extracted using the Spectrum Plant Total RNA kit from Sigma-Aldrich, and purified cDNAs were sequenced on PacBio’s Sequel II instrument.

### Scaffolding

To enable chromosome-level scaffolding of the HA412-HOv2 genome, Hi-C libraries (Burtin et al. 2013) were generated by Dovetail Genomics and sequenced to 49× depth by the McGill University and Génome Québec Innovation Centre. For the XRQv2, PSC8, IR, RHA438, PI659440, and HA89 genomes, scaffolding was aided by the production of optical maps. Briefly, ultra-HMW DNA was purified from young flash frozen leaves according to the Plant tissue DNA Isolation Base Protocol of Bionano Genomics (BNG). The ultra-HMW DNA was subsequently labelled, stained, loaded onto Saphyr chips, and run on BNG’s Saphyr platform according to the Saphyr System User Guide. Digitalized labelled DNA molecules were assembled to optical maps using BNG’s Access software.

### Genome Assembly

*De novo* assembly was conducted using different protocols depending on the genotype, the accuracy of raw sequence data and the bioinformatics tools available at the time when each genotype was sequenced (Dataset S2). In brief, the HA412-HOv2 genome was assembled with DeNovoMAGIC v3 (NRGene Technologies), and scaffolded using Hi-C sequencing data (Dovetail Genomics) and the HiRise assembler (Putnam et al. 2016).

Contigs for XRQv2, PSC8, IR, and RHA428 were generated using a meta-assembly approach (Raymond et al. 2018), whereas assembly of the other genomes used canu v2 (Koren et al. 2017). A first scaffolding step was performed for six genomes (XRQv2, PSC8, IR, RHA438, PI659440, and HA89) using BNG optical maps, and AllMaps (Tang et al. 2015) was used to anchor the sequences on the 17 chromosomes for all nine PacBio genomes.

### Genome Annotation

Gene models were predicted using the EuGene pipeline (Sallet et al. 2019), as described previously (Badouin et al. 2017). Previous RNAseq (Badouin et al. 2017) and IsoSeq (PRJNA517222) data were used for functional annotation of the HA412-HOv2, XRQv2, and PSC8 genomes. We generated IsoSeq data for the IR, RHA438, PI659440, and HA89 lines, which were employed for the annotation of each genome. IsoSeq data for HA89 were used to annotate the LR1, OQP9 and HA300 genomes. Details of the annotation processes along with assessment results generated with BUSCO v5.1.2 (-m prot -l embryophyta_odb10) software (Manni et al. 2021) are provided in Dataset S3.

To ensure that we were not over-estimating gene content variation among the ten sunflower genomes, we developed a pipeline to filter out gene fragments resulting from TE activity and other genomic processes (https://github.com/megahitokiri/Sunflower_annotation_Snakemake). At each step, parameters were fine-tuned by comparison with a set of functionally well-characterized genes to ensure the filtering was not overly aggressive. First, we employed the Extensive de novo TE Annotator (EDTA) to find areas with high content of repeated elements (Ou et al. 2019). Gene models whose exonic or 3’UTR regions overlapped more than 75% with TEs or other repetitive sequences were filtered out. The remaining gene models were further filtered to remove those with pseudogene marks, lacking introns, or that predicted proteins of less that 50 amino acids in length (Table S5).

### Identification of Sequence and Structural Variants

Because reference-guided scaffolding of the low-depth genomes (LR1, OQP8 and HA300) can cause spurious results, we only included the six high-contiguity cultivar genomes (HA412-HOv2, XRQv2, PSC8, RHA438, IR, and HA89) to identify sequence and structural variants. Each of the other five genomes was aligned to the HA412-HOv2 reference using the nucmer4 program in MUMmer v4 (Marçais et al. 2018) with parameters ‘-b 1000 -c 1000’. The alignment results were filtered using the delta-filter program in MUMmer with parameters ‘-1 -i 90 -l 1000’ to remove dubious alignments and retain only one-to-one alignments for further detection of SNPs and small InDels (<50bp). We identified SNPs and small InDels within unambiguous alignment blocks using the show-snps program in MUMmer with the parameters ‘-C -l -r -T’. The results of each pair of genomes were converted into VCF format using the HA412-HOv2 genome as the reference and the VCFs were combined using bcftools merge (Danecek et al. 2021).

We filtered the alignment results using delta-filter with parameters ‘-m -i 90 -l 1000’, and the show-coords program in MUMmer was used to extract alignment blocks with parameters ‘-T -H -r -d’ from the filtered alignment results. We then used SyRI v1.4 (Goel et al. 2019) to parse the filtered results of MUMmer to identify candidate inversions, intra-, and inter-chromosomal translocations. We merged the structural variants following a stepwise method reported in Audano et al. (2019). We set the HA412-HOv2 genome as the reference and the structural variants identified between XRQv2 and the reference as the initial callset. New sites between each genome and the reference were added in sequence. Any variants in a callset that had 50% reciprocal overlap with an existing variant was excluded. The merging was performed separately for each type of variant. Neighboring blocks belonging to same type of events were merged.

Large InDels and CNVs were identified using SVMU (Chakraborty et al. 2019) by parsing the delta file generated by delta-filter with parameters ‘-m -i 90 -l 1000’. The pipeline was run for each comparison with snp_mode = ‘l’ and without LASTZ alignments. From the SVMU summary file, structural mutations with the tag INS/DEL and estimated size >50bp were treated as large InDels (in each sample genome with respect to the HA412-HOv2 reference), and those with the tag CNV-R/CNV-Q/nCNV-R/nCNV-Q and estimated size >50bp were treated as CNVs.

### Identification of Gene Presence and Absence Variation

We constructed a pan-genome for *H. annuus* using the nine cultivated genomes plus the one wild reference sequence (Table 1). We prepared a combined GFF3/FASTA file and extracted proteins from coding regions using the TRANSDECODER (version 5.5.0-gff3_file to proteins) method (https://github.com/TransDecoder/TransDecoder). The protein files were input into the Roary pan-genome assembler (Page et al. 2015), modified to handle eukaryotic gene models, using a minimum threshold for detection of 90%, no splitting of paralogs and PRANK core genes alignment. Core alignments were assessed via a dendogram generated by Roary (SI Appendix, Fig. S22).

To distinguish between genes exhibiting true presence-absence polymorphisms and those that were annotated in one or more of the genomes but present and not annotated in others, we used representative nucleotide sequences of pan-genome genes generated by Roary to map them to each reference genome using GMAP (Wu and Watanabe 2005) with the parameters ‘-t 12 -O -n 1 -f 2 --min-trimmed-coverage=0.90 --min- identity=0.90’. Custom scripts were used to integrate the mapped genes into the pan-genome table.

### Identification of Introgressions

To identify introgressed regions in the genome assemblies of cultivated sunflower, we employed previously published resequencing data (Hübner et al. 2019; Todesco et al. 2020) from native North American landraces and five wild sunflower species (*Helianthus annuus*, *H. argophyllus*, *H. petiolaris*, *H. niveus* and *H. debilis*) that are probable donors to modern cultivated lines based on breeding records and previous studies (Vear 2016; Badouin et al. 2017; Seiler et al. 2017; Hübner et al. 2019). For each assembly, raw reads of 48 landrace and wild samples were aligned to the genome and a VCF was generated using a GATK pipeline (SI Appendix, Supplementary Information Text). Introgressed regions in the genomes were identified using the ‘site-by-site’ linkage admixture model in STRUCTURE (Pritchard et al. 2000; Falush et al. 2003).

### Projection of Introgressed Regions onto HA412-HOv2 Reference

We extracted the large alignment blocks (tag SYN/INV/TRANS/INVTR/DUP/INVDP) identified by SyRI between an assembly and the HA412-HOv2 reference as a lift-over map and converted the introgressions identified in each assembly to coordinates in the HA412-HOv2 reference. For each introgressed region, alignment blocks overlapping with the region were extracted and the positions in the original genome of the overlapping portions were projected to the reference based on the proportion relative to the start and end positions of the alignment block. Projected alignments of overlapping introgressed regions or that were within 1kb in the HA412-HOv2 reference were merged.

### Genetic Variation Analysis

The densities of SNPs and small InDels were calculated using vcftools (Danecek et al. 2011) in non-overlapping 500-kb windows. Windows overlapping with >50% with primary or secondary introgressed regions in at least one but not all genomes were defined as polymorphic introgressed windows. Densities of SNPs and small InDels were then compared between polymorphic introgressed regions and non-introgressed regions. We further annotated functional SNPs using snpEff v5.0c (Cingolani et al. 2012) and calculated the ratio of the number of alternative stop codons (*P*_nonsense_) and the number of nonsynonymous mutations (*P*_nonsyn_) in the 500-kb windows and compared polymorphic introgressed windows and non-introgressed windows within the same recombination rate category (high: > 2 cM/Mb, reduced: 0.01-2 cM/Mb, null: <0.01 cM/Mb). For the SAM population, we defined polymorphic introgressed windows as those with MAF > 0.01. SNP density and *P*_nonsense_/*P*_nonsyn_ were then calculated in non-overlapping windows of 500 kb and compared in the same way as for the genome assemblies.

For large InDels and CNVs, in each pair of genomes, we randomly sampled fragments of 500kb for 10,000 times within polymorphic primary introgressed regions, polymorphic secondary introgressed and non-introgressed regions, respectively. Densities of large InDels and CNVs were calculated and compared between these regions.

We permutated the locations of the inversions identified across the genome assemblies 10,000 times and calculated how often the overlapping size with primary introgressions and secondary introgressions exceeded the observed value, respectively. In each pair of genomes, an inversion was defined as introgression-introduced if one orientation of the inversion overlapped with primary or secondary introgressions while the other orientation did not. The incidences of inversions were calculated for polymorphic primary introgressed regions, polymorphic secondary introgressed regions and regions without introgression.

### Effects of Introgression on Gene Presence Absence Variation

To determine how introgression affected gene content, we filtered the table of gene presence-absence polymorphism based on synteny between the genomes as determined by MUMmer4 (Marçais et al. 2018). Using the synteny-filtered table of gene presence-absence polymorphisms, as well as the introgressions identified in each genome, we assigned a single introgression value for each gene in a genome if > 50% of the gene overlapped with regions of primary or secondary introgressions. Each missing copy in a genome was assigned an introgression value if the corresponding MUMmer alignment overlapped >50% with regions of primary or secondary introgressions. We compared each of the cultivar genomes to the HA412-HOv2 reference and examined the presence/absence of genes in introgressed and non-introgressed regions.

### Effects of Introgressions on Phenotypic Variation in the SAM Population

We made use of 287 cultivated accessions in the SAM population, which was previously sequenced to 5-25x depth (Hubner et al. 2019). The SAM population includes close to 90% of cultivated sunflower genetic diversity (Mandel et al. 2011) and is comprised of both inbred and open-pollinated lines, as well as oilseed and confectionary cultivars. All 287 accessions, as well as the aforementioned 48 landrace and wild samples, were mapped to the HA412-HOv2 reference genome, and a SNP data set was generated using a pipeline similar to that described above (SI Appendix, Supplementary Information Text). We then used the SNP data set to identify introgressions from the primary and secondary germplasm in all accessions using the software package PCAdmix (Brisbin et al. 2013), a principal component analysis-based algorithm for inferring local ancestry along chromosomes in admixed genomes. Prior to the PCAdmix analysis, the VCF was filtered to retain only bi-allelic SNPs in the 50% tranche from GATK Variant Quality Score Recalibration with genotyping rate > 90%, and the SNPs were phased using Beagle 5.1 (Browning et al. 2018) for each species separately. No pruning was set in the PCAdmix analyses.

The identified introgressed regions from wild *annuus* and secondary germplasm were used to call introgression variants in the SAM population. We assessed the presence or absence of introgressions in 1kb non-overlapping windows across the genome of each sample in the SAM population. Introgression variants were subsequently filtered for minor allele frequency ≥ 3%.

For the phenotypic analyses, we employed data for 16 traits that were generated as part of a common garden study carried out in 2011 at three locations: Watkinsville, GA and Ames, IA in the USA and Vancouver, BC, in Canada (Mandel et al. 2013; Nambeeson et al. 2015; Lee et al. 2022). To identify associations between introgression variants and the phenotypic traits, a genome wide association (GWA) analysis was carried out using EMMAX (Kang et al. 2010). Population structure was corrected by the first three principal components of the LD-pruned SNP dataset (calculated with PLINK --indep-pairphase 50kb 50 0.2; Purcell et al. 2007). To correct for relatedness between samples in the GWA analysis, the SNP dataset was used to estimate a kinship matrix by EMMAX.

To identify significantly associated introgression markers and the direction of the introgression on phenotypic data, we generated double-sided Manhattan plots, in which introgression markers that increase or decrease trait values were shown with − log10(P-value) and log10(P-value), respectively. To avoid false-positive associations, Bonferroni correction was used as the threshold of significant association.

To further explore the signature of linkage drag on phenotypic data, a ridge regression model was used to estimate the effect of each introgression variant on a given trait with the mixed.solve function in R package rrBLUP version 4.6.1 (Endelman et al. 2011). The average effect size of introgressions for each trait was compared to a null distribution.

We assessed the significance of an introgression variant’s effect on phenotype variation by testing whether the observed impact size of introgressions was either larger than the 95th percentile of the tail of the null distribution (significantly larger) or smaller than the 5th percentile of the tail of the null distribution (significantly smaller). To construct the null distribution, 10,000 introgression effect size estimates for each trait were generated by shuffling introgression variants between samples and calculating the average effect size of introgressions. We further compared the effect size of introgression on each trait for primary versus secondary germplasm donors.

A linear model (Y ∼ X) was fit to evaluate the effects of frequency on the phenotypic impact of introgression, where Y is a vector of introgression effect and X is a vector of introgression frequency. The beta coefficient of X can therefore represent the contribution of frequency to the direction and effect size of introgression variants.

### Data Availability

Genome assemblies and annotations are available at https://www.heliagene.org/ and https://sunflowergenome.org/ for the PacBio and Illumina genomes, respectively. Raw sequences are deposited in NCBI (Table S8). Custom scripts for the analyses are available upon request and will be sent to GitHub before publication.

## ACKNOWLEDGMENTS

We thank NRGene and DoveTail Genomics for assembly and scaffolding, respectively, of the HA412-HOv2 genome, Greg Baute for comments and discussions during the project. This work was supported by the International Consortium of Sunflower Genomics, a China Scholarship Council scholarship (no. 201506380099) to KH, a grant from the NSF Plant Genome Program (IOS-1444522 to JMB and LHR), and a National Science and Engineering Discovery Grant to LHR. We are grateful to Compute Canada and the GenoToul bioinformatics platform of Toulouse-Occitanie for providing computing and storage resources.

