## Supplementary material for "The genomics of linkage drag in sunflower": SI Appendix, Fig. S1

### Supplementary Information Text

### Material and Methods

#### Diversity Analyses

To illustrate the genetic positioning of the nine sequenced cultivars in the cultivated sunflower gene pool (Table S1), we analyzed an unpublished data set comprising 16,048 SNP markers (AXIOM SUNAX51, Mangin et al., 2017) and 2,850 cultivated lines. These include 2,606 lines from France's National Research Institute for Agriculture, Food and Environment (INRAE) genebank, 134 lines from the SAM population, which are United States Department of Agriculture (USDA) genebank lines, 99 lines from Argentina's National Agricultural Technology Institute (INTA) core collection, and 11 lines from the Inner Mongolia University core-collection. SNP markers were selected if they were homozygous and constant on five replicates of the XRQ genotype and missing data imputed as the XRQ value. Principle Components Analysis (PCA) was performed using the PCA function from FactoMineR R package (Lê et al., 2008) with five dimensions to construct the genetic space presented in Fig. S1.

#### Nucleic Acid Extractions, Library Preparations, and Sequencing

Seeds of HA412-HO were surface-sterilized by immersion in a 20% bleach solution for 10 minutes, rinsed with distilled water, and directly sown in Sunshine potting mix #1 (Sun Gro Horticulture). Soon after emergence, seedlings were put in the dark for 4 days, and etiolated young leaves were collected and flash-frozen in liquid nitrogen. DNA was extracted using a modified CTAB protocol, as in Todesco et al. 2020.

For the other cultivated accessions, three-week-old plantlets were harvested, frozen in liquid nitrogen, and stored at -80°C prior to manual crushing with pre-cooled mortar and pestle in liquid nitrogen. High molecular weight DNA was subsequently isolated following the magnetic bead protocol of Mayjonade et al. (2016). In brief, a buffer containing SDS was used to lyse cells walls, followed by the addition of RNase A to inhibit RNA activity. Next, proteins and polysaccharides were removed using a potassium acetate buffer and a phenol/chloroform mixture, followed by centrifugation. DNA was subsequently purified using carboxylated magnetic beads (SpeedBeads magnetic carboxylate modified particles, GE Healthcare), followed by two wash steps with 70% ethanol and elution in a buffered solution. DNA purity, quantity, and size were measured via UV spectrophotometry, Qubit fluorometry (Thermo Fisher Scientific), and the Femto Pulse System (Agilent Technologies), respectively (Table S2).

Seeds of the wild *H. annuus* accession PI659440 (Nebraska, USA) were obtained from the USDA GRIN system (https://www.ars-grin.gov/), grown in 3L pots in a controlled growth room, and selfed for three generations to reduce heterozygosity. Young leaves from third generation plants were harvested in Israel and sent to France, where they were stored at -80°C prior to DNA extractions. High molecular weight DNA extracted using magnetic beads (Mayjonade et al. 2016) was of poor quality, so the QIAGEN Genomic-tip 100g procedure was employed instead. DNA purity, quantity, and size were estimated as described above (Table S2).

For RNA extractions, leaves, flowering buds and stems of IR, RHA438, PI659440, and HA89 lines were harvested from sequenced plants, frozen in liquid nitrogen, and stored at -80°C. Leaves were sampled from 3-week-old plants, whereas buds and stems were obtained from 3-month-old plants. RNA extraction was done using the Spectrum Plant Total RNA kit (Sigma-Aldrich), followed by on-column DNase Digestion (DNase10) to remove all traces of DNA. Quantity and quality of RNA was assessed using Agilent Technologies' 2100 Bioanalyzer system (Table S3).

**Library Preparations and Sequencing**

For the HA412-HOv2 genome, PCR-free paired-end (250bp×2 and 150bp×2) and Nextera™ MP Gel Plus mate-pair (150bp×2) libraries were developed with insert sizes ranging from 450 bp to10 kb and sequenced using Illumina sequencing technology to a total depth of 214× (Dataset S1). In addition, 10× Genomics Chromium libraries (150bp×2) were prepared and sequenced using Illumina technology to 37× depth (Dataset S1).

The remaining nine genotypes were sequenced using Pacific Biosciences (PacBio) sequencing technology at the Gentyane Sequencing Platform (Clermont-Ferrand, France). Data were generated between 2015 and 2021 depending on the genotype (Dataset S1). For XRQv2 and PSC8, sequencing libraries were prepared following PacBio's "Procedure & Checklist - 20 kb Template Preparation". HMW-DNA was sheared at 40kb using the Megaruptor system (Diagenode), followed by an 18-kb size selection using the BluePippin system (Sage Science). The libraries were sequenced using P6/C4 chemistry with 6-hour movies on a PacBio RSII instrument.

For IR and RHA438, CLR (Circular Long Reads) libraries were prepared using the SMRTbell Express 2 Template prep kit, following PacBio's "Procedure and Checklist - Preparing gDNA Libraries using SMRTbell Express Template prep kit 2.0" protocol. High molecular weight genomic DNA (12 ug) was sheared with the 75 kb program using a Diagenode Megaruptor, which generated DNA fragments of ca. 60 kb. A Femto Pulse assay was used to assess fragment size distributions. Sheared genomic DNA were carried into the enzymatic reactions to remove the single-strand overhangs and to repair any damage that may be present on the DNA backbone. An A-tailing reaction followed by overhang adapter ligation was conducted to generate the SMRTBell template. After purification with 0.45X AMPure PB beads, size-selection was performed using the BluePippin to recover fragments of >30 kb, leading to final libraries in the 40 kb range. Quality and quantity of the SMRTBell libraries were determined using the Femto Pulse System and a Qubit Fluorimeter with Qubit dsDNA HS reagent Assay kit (Life Technologies). Ready-to-sequence SMRTBell Polymerase Complexes were created using the Sequel II Binding Kit 2.0 (PacBio) and the v4 primer. The PacBio Sequel II instrument was programmed to load a 45 pM library and sequenced on a Sequel II Sequencing plate v2.0 with one movie of 30 hours per SMRTcell, using the CLR mode.

For IR, RHA438, HA89, HA300, LR1, OQP8 and PI659440, high-fidelity (HiFi) libraries were produced using SMRTbell Express 2 Template prep kit, following PacBio's "Procedure and Checklist - Preparing HiFi SMRTBell Libraries using SMRTbell Express Template prep kit 2.0" protocol. High molecular weight genomic DNA (15 ug) was sheared with the Diagenode Megaruptor as above, except the 25 kb program was used, yielding DNA fragments of ca. 20 kb. The remainder of the protocol was similar to that described for IR and RHA438, except that 1X AMPure PB beads were used for DNA purification, and the BluePippin system was employed to recover DNA fragments > 10 kb, resulting in final libraries of ca. 20 kb. Ready-to-sequence SMRTBell Polymerase Complexes were created using the Sequel II Binding Kit 2.0 (PacBio) and the v2 primer for IR, RHA438, PI659440 or the Sequel II Binding Kit 2.2 and the v5 primer for OQP8, HA300 and LR1. The Sequel II was programmed to load an 80 pM library and samples were sequenced on a Sequel II Sequencing plate v2.0 with 2 hours of pre-extension and one movie of 30 hours per SMRTcell, using CCS mode.

For IR, RHA438, HA89 and PI659440, Iso-Seq SMRTbell libraries were constructed according to the standard isoform sequencing protocol "Procedure and Checklist-Iso-Seq ™ Express Template Preparation for Sequel® and Sequel II Systems" using the NEBNext Single Cell/Low Input cDNA Synthesis & Amplification Module (New England Biolabs) and the ProNex Size-Selective Purification System (Promega) for size selection. Briefly, 300 ng of total RNA from different tissues (leaves, stems, flower buds) was used as input for complementary DNA synthesis. Each sample was barcoded and then subjected to cDNA amplification using 12 cycles. Purified cDNAs were pooled in equal molarity and subjected to library preparation using PacBio's SMRTbell Express Template Prep Kit 2.0 following the Iso-Seq protocol previously referenced. The library was prepared for sequencing by annealing primer v4 with the Sequel II Binding Kit 2.1. The PacBio Sequel II instrument was programmed to load a 70 pM library and the sample was sequencing on a Sequel II Sequencing plate v2.0 with one movie of 24 hours per SMRTcell.

**Optical Mapping**

To generate optical maps, ultra-HMW DNA was purified from 0.5 g of young flash frozen leaves according to the Plant tissue DNA Isolation Base Protocol of Bionano Genomics (BNG). Leaves were ground in liquid nitrogen and then homogenized in a homogenization buffer (BNG) using a TissueRuptor grinder (Qiagen). Nuclei were washed and embedded in agarose plugs. After overnight proteinase K digestion in Lysis Buffer (BNG) and 1-hour treatment with RNAse A (Qiagen), plugs were washed 4 times in 1×Wash Buffer (BNG) and 5 times in 1× TE Buffer (ThermoFisher Scientific). Then, plugs were melted for 2 minutes at 70 °C and solubilized with 2 µL of 0.5 U/µL AGARase enzyme (ThermoFisher Scientific) for 45 minutes at 43 °C. A dialysis step was performed in 1× TE Buffer (ThermoFisher Scientific) for 45 minutes to purify DNA from any residues. The DNA samples were quantified using the Qubit dsDNA BR Assay (Invitrogen). Quality of megabase-size DNA was validated by pulsed-field gel electrophoresis.

Labelling and staining of the ultra-HMW DNA were performed according to the Direct Label and Stain protocol (BNG). Labelling was performed by incubating 750 ng genomic DNA with 1× DLE-1 Enzyme (BNG) for 2 hours in the presence of 1× DL-Green (BNG) and 1× DLE-1 Buffer (BNG). Following proteinase K digestion and DL-Green clean-up, the DNA backbone was stained by mixing the labelled DNA with DNA Stain solution (BNG) in the presence of 1×Flow Buffer (BNG) and 1× DTT (BNG), and incubated overnight at room temperature. The DLS DNA concentration was measured with the Qubit dsDNA HS Assay (Invitrogen). Labelled and stained DNA was loaded on Saphyr chips (BNG). Loading of the chips and running of the BNG Saphyr System were performed according to the Saphyr System User Guide. Digitalized labelled DNA molecules were assembled into optical maps using the BNG Access software.

**Genome Assembly**

For the HA412-HOv2 genome, sequencing data were processed and assembled by NRGene using their DeNovoMAGIC assembler v3.0. To enable chromosome-level scaffolding, Hi-C libraries (Burtin et al. 2013) were generated by Dovetail Genomics and sequenced to 49× depth by the McGill University and Génome Québec Innovation Centre. The Hi-C data were subsequently employed by Dovetail Genomics to further improve and scaffold the DeNovoMAGIC assembly with the HiRise software platform (Putnam et al. 2016).

XRQv2 (updated version of previously published XRQ genome, Badouin et al. 2017) and the remaining eight genotypes were sequenced using PacBio sequencing technology (Table 1). Data were generated between 2015 and 2021 depending on the genotype (Dataset S1). Consequently, genome assembly protocols differ depending on the accuracy of raw sequence data and the evolution of bioinformatics software. Dataset S2 describes the main steps of the nine PacBio genome assemblies, the evolution of the metrics during the assembly process, and the software versions.

Contigs for XRQv2, PSC8, IR, and RHA438 were generated using a meta-assembly approach based on the protocol and til-r software developed to assemble the *Rosa* genome (Raymond et al. (2018). The rationale was to assemble the results of several primary assemblies using different software with different heuristics and parameters to increase contiguity and minimize gaps in the final assembly. More recent and accurate Sequel II HiFi reads were generated for PI659440, HA89, LR1, OQP8 and HA300 (Dataset S1). The increased accuracy provided by HiFi reads, combined with improvements in the canu v2 software (Koren et al. 2017), made the meta-assembly strategy obsolete. Canu v2 integrates additional heuristics to remove spurious edges in the overlap graph, making it less sensitive to parameter changes when using HiFi reads. Consequently, a single assembly was performed to generate the contigs of the PI659440, HA89, OPQ8, LR1, and HA300 genotypes. Spurious contigs – hits with a larger contig spanning 80% of the length of the shorter contig – were removed prior to scaffolding using minimap2 (-x asm5) software (Li 2018).

A first scaffolding step was performed for six genomes (XRQv2, PSC8, IR, RHA438, PI659440, and HA89) using Bionano optical maps (https://bionanogenomics.com/downloads/bionano-solve/). When a version of the Bionano Solve anterior to 3.5 was used, a post process based on ncbi-blastn was applied to close false gaps due to undetected overlaps (issue corrected since version 3.5).

For all nine PacBio genomes, AllMaps (Tang et al. 2015) was used to anchor the sequences onto the 17 sunflower chromosomes. XRQv2 and PSC8 were anchored using three genetic maps described in Badouin et al. 2017) to build the XRQv1 genome. IR, RHA438, PI659440, and HA89 were anchored using the XRQv2, PSC8, and HA412-HOv1 genomes. The three reference genomes were first repeat-masked using Red (-len 16) software (Girgis 2015), then mapped on the genomes to be anchored with minimap2 (-x asm5). Hits longer than 10kb were converted in genetic markers in the format expected by AllMaps (weight set to 1 for all genomes). LR1, HA300, and OPQ8, for which the sequencing depth is low and no optical maps available, were anchored using XRQv2, PSC8, HA412-HOv1, PI659440, and HA89 genomes as references.

The pseudomolecule sequences of XRQv2 and PSC8 were polished twice. First using PacBio ARROW software (https://www.pacb.com/support/software-downloads/), and then with illumina paired-end data and Pilon software (Walker et al. 2014).

**Annotation of Transposable Elements**

We conducted de-novo annotation of transposable elements (TEs) for the genome assemblies using the program EDTA (Ou et al. 2019). The program was run with the parameters "--sensitive 0 --evaluate 0 --anno 1" to perform identify raw TE candidates, filter out false discoveries and generate whole-genome TE annotation on the 17 chromosomes of each genome assembly.

**Centromere Locations**

We mapped the locations of centromeres in the genome assemblies using the sequences previously found to be targeted by sunflower centromere-specific histone H3 (Nagaki et al. 2015; Huang et al. 2022). The main sequence from Nagaki et al. (2015) was downloaded from the Nucleotide database of National Center for Biotechnology Information (NCBI) with the accession number LC075745, and the sequence was then queried against each genome using BLASTN (https://blast.ncbi.nlm.nih.gov/) with an E-value of $1\times10^{-5}$.

**Identification of Introgression**

For each assembly, raw sequencing data from 16 landrace individuals and 10 samples with highest sequencing depths from each of the five wild species were clipped and filtered to remove Illumina adapters, poor quality bases and short (<36 bp) reads using Trimmomatic v0.36 (Bolger et al. 2014), and the filtered reads were aligned to the genome using Next-GenMap v.0.5.3 (Sedlazeck et al. 2013). The resulting BAM files were concatenated, sorted, and duplicate-marked using samtools v0.1.19 (Li et al. 2009). Libraries sequenced in multiple lanes were merged with sambamba v0.6.6 (Tarasov et al. 2015), and PCR duplicates were marked with samtools. Variant calling was performed following best practices of Genome Analysis Tool Kit v 4.1.4.1 (GATK; McKenna et al. 2010). For each sample, a GVCF file was produced with the GATK 'HaplotypeCaller' with the parameter "--heterozygosity 0.01". After individual variant calling, all samples were jointly genotyped using GATK's 'GenomicsDBImport' and 'GenotypeGVCFs'. To reduce computational time, genomic regions containing TEs and small unplaced contigs were excluded, and the step was run over 1 Mb regions for parallel computation. The raw VCF chunks were then gathered by chromosome using 'GatherVcfs'. The 10 samples with the highest sequencing depth were selected to produce a "gold set" using the following parameters: mapping quality > 50.0, missing rate < 10%, -1.0 < strand odds ratio < 1.0, minor allele frequency > 0.25, excess heterozygosity < 10.0, -1.0 < BaseQRankSum < 1.0, depth within one standard deviation from the mean and ExcessHet z-score > -4.5. The raw set of all variants were first filtered to remove sites with extremely heterozygosity (ExcessHet z-score < -4.5) and the gold set was then applied against this filtered set of variants to produce recalibration models using 'VariantRecalibrator'. Variants in the 90% tranche was selected using 'ApplyVQSR' and subsequently filtered to retain only bi-allelic SNPs with minor allele frequency > 0.01 and genotyping rate > 50%.

We ran principal component analysis (PCA) using the R package "SNPRelate" (Zheng et al. 2012) and constructed Neighbor-joining trees using MEGA 10.1.6 (Kumar et al. 2018) with the maximum composite likelihood method to examine relationships between samples. We filtered 2 samples of *H. debilis* that displayed abnormal clustering pattern and selected 8 individuals with highest sequencing depths from landraces and each of the other wild species to ensure balanced sampling. The VCF of the remaining 48 samples was filtered to retain only bi-allelic SNPs in the 50% tranche with minor allele frequency > 0.05 and genotyping rate > 80%.

To identify introgressed regions in the genomes, we used the 'site-by-site' linkage admixture model in STRUCTURE (Pritchard et al. 2000). The genotypes were formatted for STRUCTURE using custom Perl scripts. Genotype calls of each genome assembly at each site were taken from the reference sequence. STRUCTURE was run separately on each chromosome using the following parameters: LOG10RSTART = -6.124939, LOG10RMIN = -9, LOG10RMAX = -5.823909, LOG10RPROPSD 0.1, ADMBURNIN = 500, BURNIN = 10000, NUMREPS = 10000. Since modern lines were largely derived from landrace *H. annuus*, sites with ancestry (> 0.95) for a non-landrace group were defined as introgressed (Baute et al. 2015, Badouin et al. 2017). Introgressed regions in each genome were extracted by dividing the genome in windows with divisions at the midpoint between SNPs (Badouin et al. 2017), and adjacent introgressed regions from the same non-landrace group were merged.

**Genetic Variation Analysis**

We compared SNPs and small InDels between regions with and without introgression by a window-based analysis. We divided the reference (HA412-HOv2) into non-overlapping 500-kb windows, and for each genome, windows with >50% overlap with the projected introgressed regions were defined as introgressed. Across the six high-contiguity cultivar genomes, windows introgressed in at least one but not all genomes were defined as polymorphic introgressed windows. Introgressions from the primary gene pool and those from the secondary gene pool were assessed separately. The densities of SNPs and small InDels on the 17 chromosomes were calculated using vcftools (Danecek et al. 2011) with a window size of 500kb. Windows with less than 50% coverage by one-to-one MUMmer alignments were removed due to lack of information. The densities of SNPs and small InDels were then compared among introgressed and non-introgressed regions. We analyzed primary introgressions and secondary introgressions separately. The number of SNPs and small InDels were log-transformed and compared using Welch two sample t-tests.

We further annotated the VCFs using the program snpEff v5.0c (Cingolani et al. 2012) with protein-coding genes from the HA412-HOv2 annotation. Synonymous mutations ('synonymous_variant'), nonsynonymous mutations ('missense_variant') and alternative stop codons ('stop_gained' and 'stop_lost') were extracted from the annotated VCFs. The deleterious load was represented as the ratio of the number of alternative stop codons ($P_{nonsense}$) and the number of nonsynonymous mutations ($P_{nonsyn}$) (Renaut and Rieseberg 2015). We calculated this statistic in sliding windows of 500 kb and compared it to recombination rate and among introgressed and non-introgressed regions. To control for the effect of recombination, we divided the windows into three different categories based on recombination rate: (1) high recombination rate for windows with recombination rate > 2 cM/Mb, (2) reduced recombination rate regions with recombination rate between 0.01 and 2 cM/Mb, and (3) null recombination regions with recombination rate < 0.01 cM/Mb (Bartolome et al. 2002). Polymorphic introgressed windows were compared to non-introgressed windows of the same recombination rate category. Primary and secondary introgressions were analyzed separately. $P_{nonsense}/P_{nonsyn}$ was logit-transformed and compared using Welch two sample t-tests.

We also examined the effect of introgression on SNP density as well as genetic load using the data sets generated using resequencing data of 287 cultivated accessions in SAM population (see below). We defined polymorphic introgressed windows in the SAM population as those with MAF > 0.01 (primary introgressions and secondary introgressions separately) and used windows that did not overlap with primary introgressions or secondary introgressions as a control. We then calculated SNP density and $P_{nonsense}/P_{nonsyn}$ in sliding windows of 500 kb and compared it to recombination rates and local ancestry in the same way as for the genome assemblies.

In each high-contiguity genome mapped to the HA412-HOv2 reference, we compared large InDels and CNVs between polymorphic introgressed regions and non-introgressed regions for primary introgression and secondary introgression, respectively. For each type of region, we randomly sampled fragment of 500kb within the regions and calculated the densities of large InDels and CNVs within the sampled fragment. We repeated this process 10,000 times and compared the observations of non-introgressed, polymorphic primary introgressed and polymorphic secondary introgressed regions using Welch two sample t-tests.

For inversions identified across the six high-contiguity genome assemblies, we assessed the overlap of the inversions with introgressions. We first estimated an empirical null distribution for the overlap of inversions with polymorphic introgressed regions by permutating the inversions across all chromosomes 10,000 times and then asked how often the overlap with primary introgressions and secondary introgressions exceeds the observed value; a one-tailed p-value was calculated to determine significance. For each pairing of a cultivar genome and the HA412-HOv2 reference, we defined an inversion as introgression-introduced if one orientation of the inversion overlapped with primary or secondary introgressions in the genome in which it resided while the other orientation did not. The number of inversions introduced by primary and secondary introgressions were normalized by the total size of polymorphic introgressions between genomes.

**Effects of Introgression on Gene Presence Absence Variation**

To study the effect of introgression on the presence/absence of genes, we filtered the table of gene presence-absence polymorphism based on synteny between the genomes. We performed pairwise alignments using nucmer in MUMmer4 (Marçais et al. 2018) with the parameters '-mum -c 1000', filtered the alignments using delta-filter with the parameters '-g -u 70', and created tables to identify the syntenic positions with show-cords using the parameters '-c -l -r -T'. MUMmer alignments that covered >40% of each of Roary's representative gene were extracted and the corresponding alignment in each genome were checked. We kept the homologs of each gene if a homolog was within the upper or lower 10kb of the alignment. If a genome did not have a homolog of the representative gene or the genes did not pass the synteny filter, we defined the gene as missing in the genome and kept the location of the alignment for downstream analyses. When no MUMmer alignments were found between genomes that covered a Roary's representative gene, we identified the lower and upper boundaries of the closest alignments and used the overlapping of the gap regions and the representative gene to determine the presence/absence of genes using the same method as above. Cytoplasmic genes were treated as missing, and genes that did not have synteny information or those on unanchored contigs were excluded from the analysis.

Using the synteny-filtered table of gene presence-absence polymorphisms, as well as the introgressions identified in each genome, we assigned a single introgression value for each gene in a genome if > 50% of the gene overlapped with regions of primary or secondary introgressions. Each missing copy in a genome was assigned an introgression value if the corresponding MUMmer alignment overlapped >50% with regions of primary or secondary introgressions. We compared each of the eight cultivar genomes to the HA412-HOv2 reference and examined the presence/absence of genes in introgressed and non-introgressed regions.

**Identification of Introgressions in the Cultivated Sunflower Association Mapping (SAM) Population**

All the 287 cultivated accessions in the SAM population, as well as the aforementioned 66 landrace and wild samples, were mapped to HA412-HOv2 reference genome, and variant calling was performed used a similar pipeline described above. For variant recalibration, 20 samples in the SAM population with the highest sequencing depth were chosen to produce the "gold set". The variants were subsequently filtered to retain only bi-allelic SNPs in the 90% tranche with minor allele frequency > 0.01 and genotyping rate > 50%.

We used the SNP dataset called across wild samples, landraces, and all accessions to identify introgressions using the software package PCAdmix (Brisbin et al. 2012). As in the identification of introgression in the reference genomes, we discarded samples with abnormal clustering pattern and selected 8 individuals with highest sequencing depths from landraces and each of the wild species. A VCF with the 48 samples and samples in the SAM population was generated and filtered to retain only bi-allelic SNPs in the 50% tranche with genotyping rate > 90%. The SNPs were then phased for each species with a linkage map using Beagle 5.1 (Browning et al. 2018), and the genotype tables were then formatted for PCAdmix using the vcf2beagle program in Beagle 5.1. No pruning was set within the PCAdmix analyses (-prune 0).

We first pooled all wild samples to form a single ancestral group and assigned the landrace individuals to another ancestral group as the most probable source of variation. PCA loadings were computed by PCAdmix for the ancestral haplotypes and for the individuals of the SAM population by treating them as admixed individuals, and the genome-wide and local ancestries of the admixed individuals was inferred from the PCA using a window size of 100 SNPs (-w 100). The signal was then smoothed using the Viterbi algorithm within an HMM framework in PCAdmix (Brisbin et al. 2012). Posterior probabilities of ancestry affiliation at every window in the admixed individuals were obtained from the HMM, and the most likely ancestry classification at each window was extracted to identify introgression from wild species. Introgressed regions were defined with divisions at the midpoint between the first SNP in a window and the last SNP in the previous window. Adjacent introgressed regions were merged. We later pooled samples from *H. petiolaris*, *H. niveus* and *H. debilis* into a single ancestral group (PET clade) according to the results of PCA and phylogeny of the samples (Fig. S23) and ran PCAdmix using the ancestral combination of *H. annuus*, *H. argophyllus* and the PET clade as well as the ancestral combination of landraces, *H. argophyllus*, and the PET clade using the same settings as the analysis above. Introgressions from each ancestral group were identified with the same algorithm, and adjacent introgressed regions from the same ancestral group were merged. The introgressions from *H. argophyllus* and the introgressions from the PET clade identified in both runs were treated as trustable introgressions from the secondary gene pool. The introgressions from wild species that did not overlap with introgressions from secondary gene pool were assumed to derive from the primary gene pool.

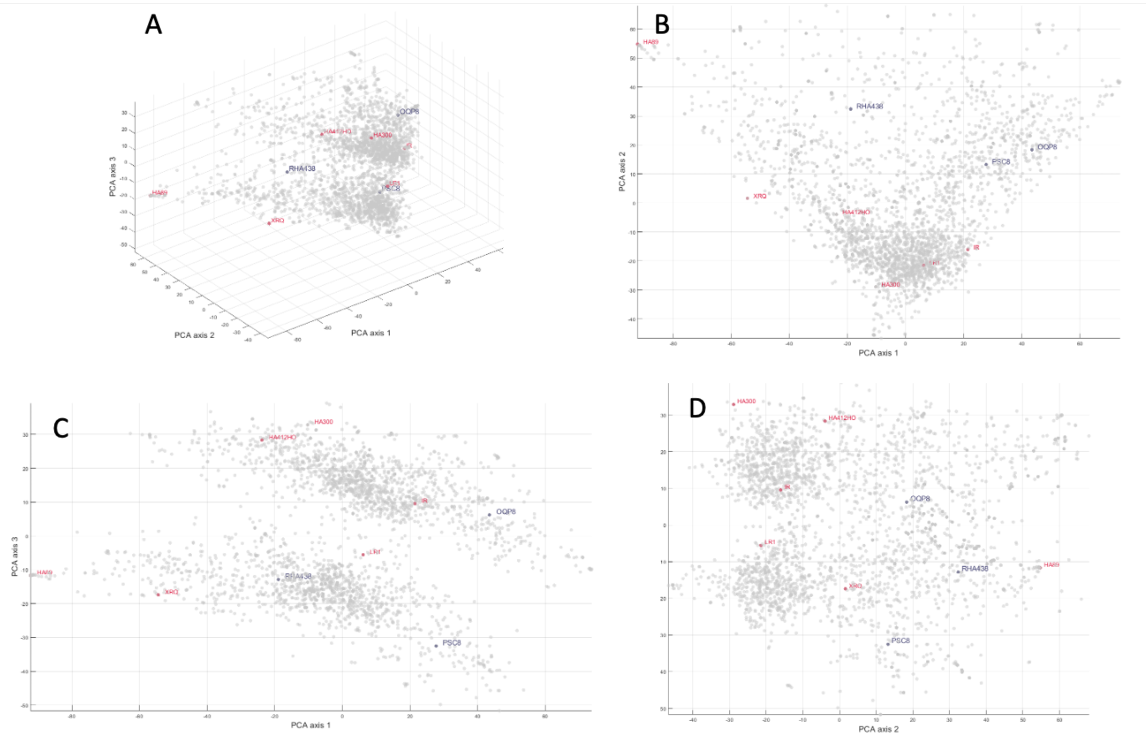


**Figure S1.** Sequenced lines in the genetic space constructed by Principal Component Analysis (PCA) from 16,048 SNP markers and 2,716 inbred lines present in INRAE, USDA, INTA, and University of Inner Mongolia genebanks. (A) 3D view, (B) PCA axis 1 and 2, (C) PCA axis 1 and 3, (D) PCA axis 2 and 3. In blue restorer lines of cmsPET1, in red maintainer lines of csmsPET1.

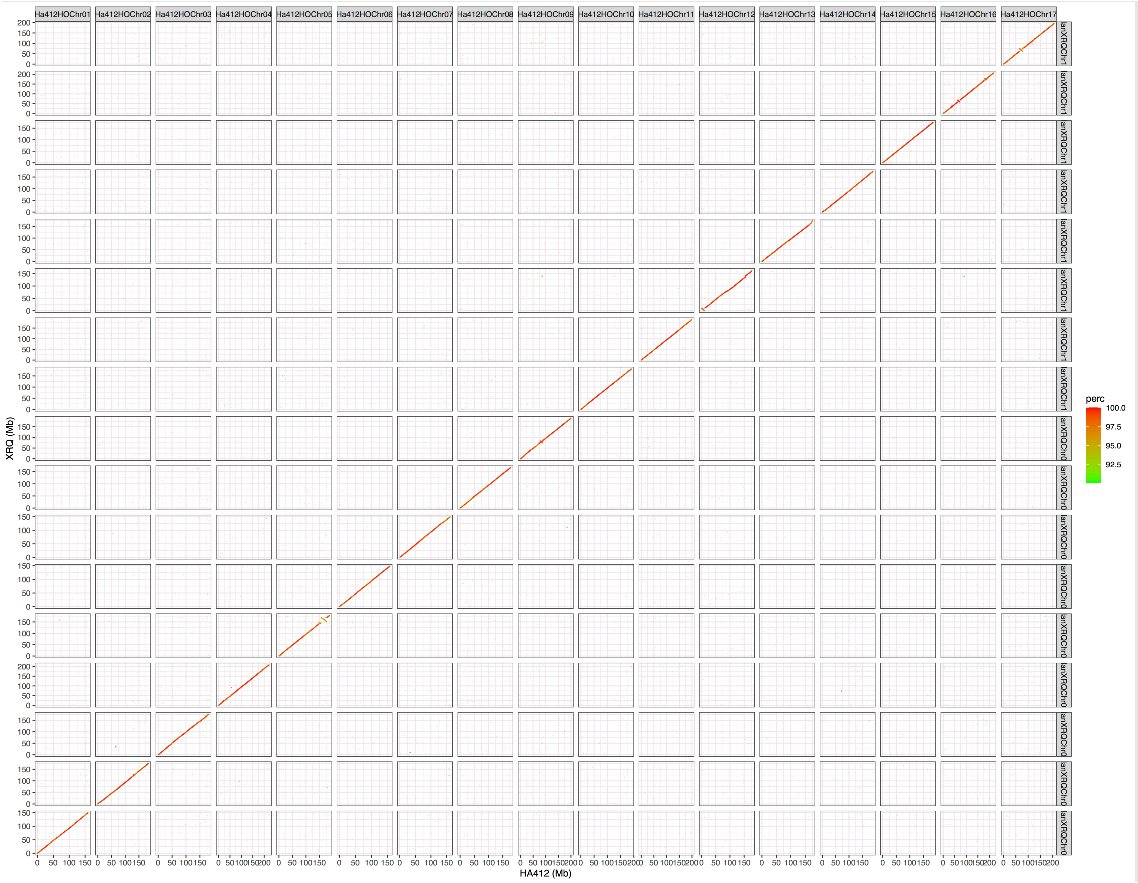


**Figure S2.** MUMmer alignment between XRQv2 and the HA412-HOv2 reference.

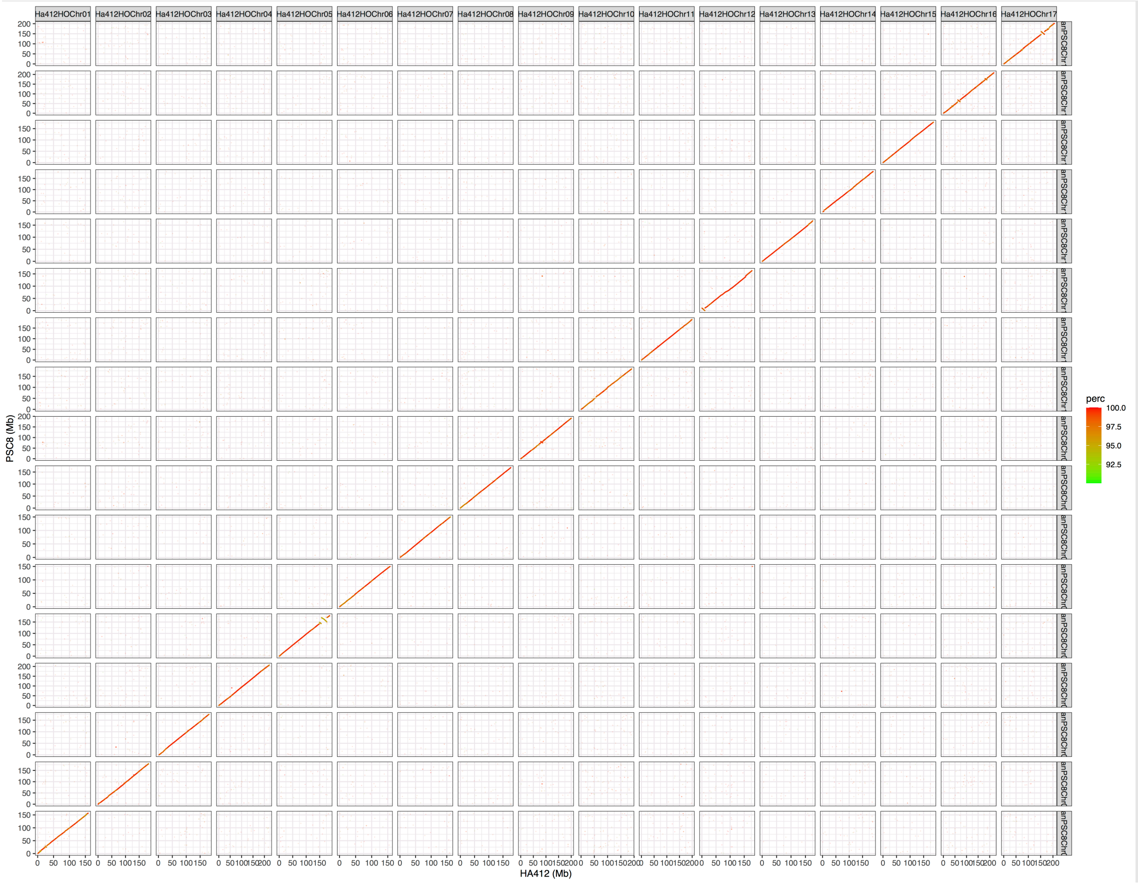


**Figure S3.** MUMmer alignment between PSC8 and the HA412-HOv2 reference.

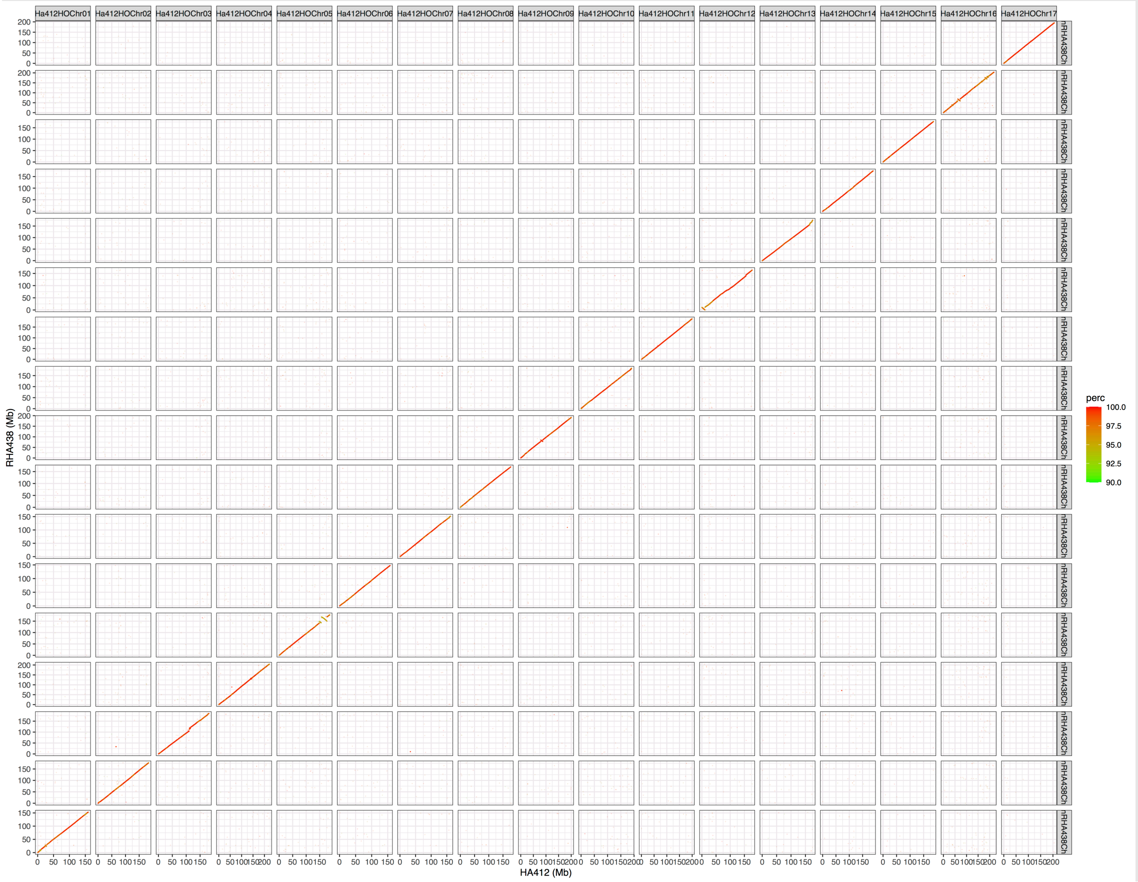


**Figure S4.** MUMmer alignment between RHA438 and the HA412-HOv2 reference.

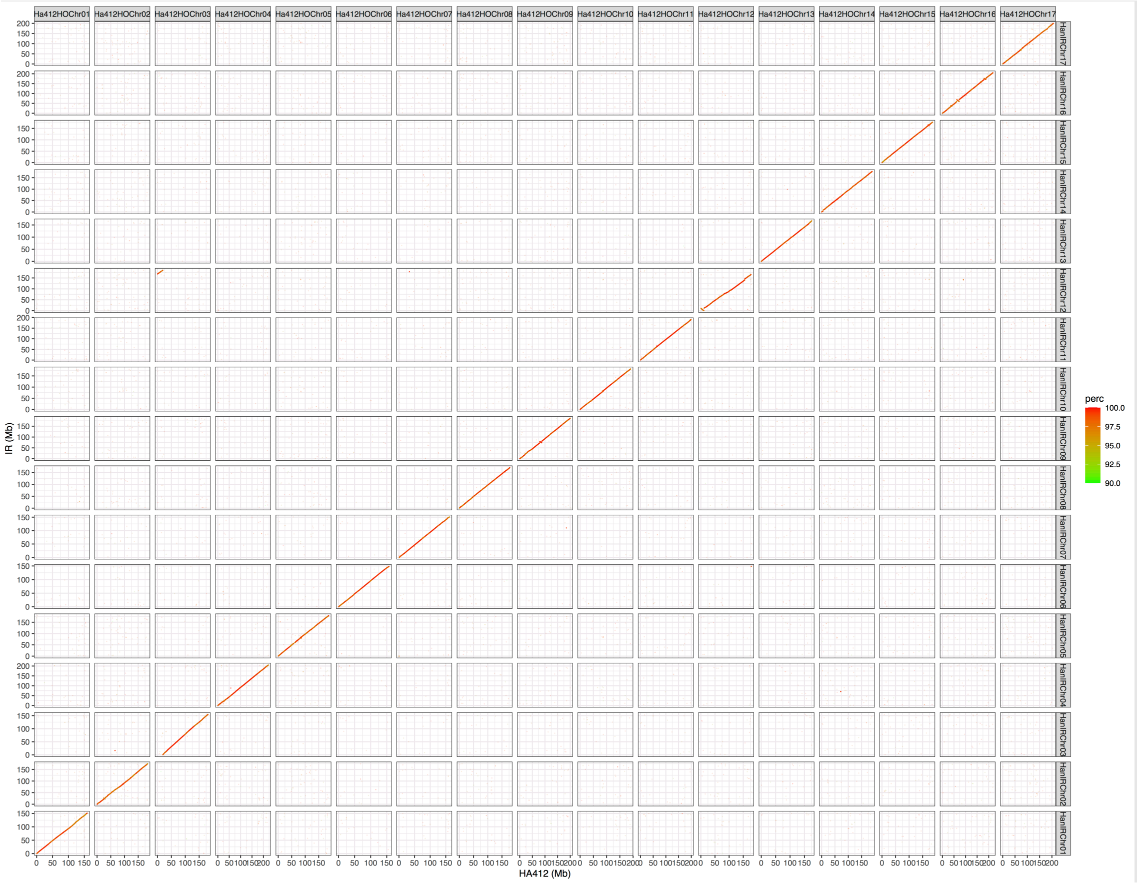


**Figure S5.** MUMmer alignment between IR and the HA412-HOv2 reference.

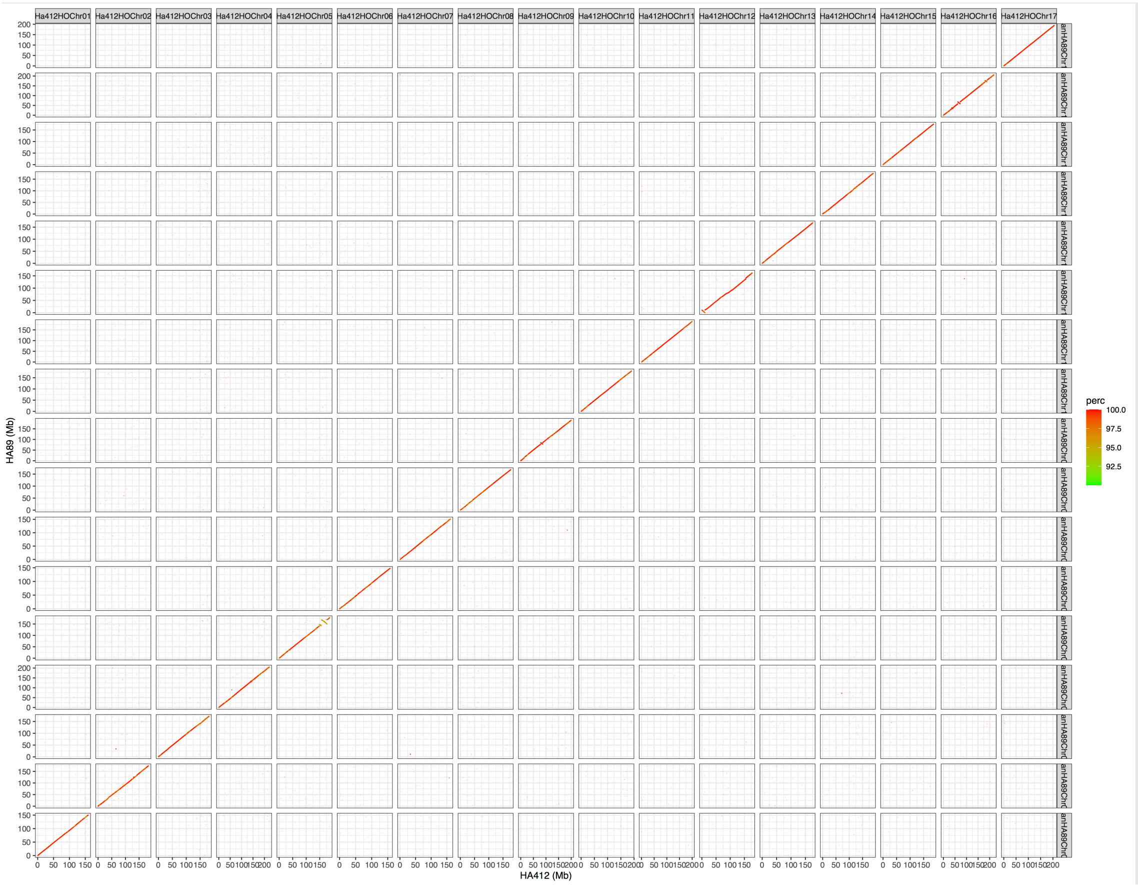


**Figure S6.** MUMmer alignment between HA89 and the HA412-HOv2 reference.

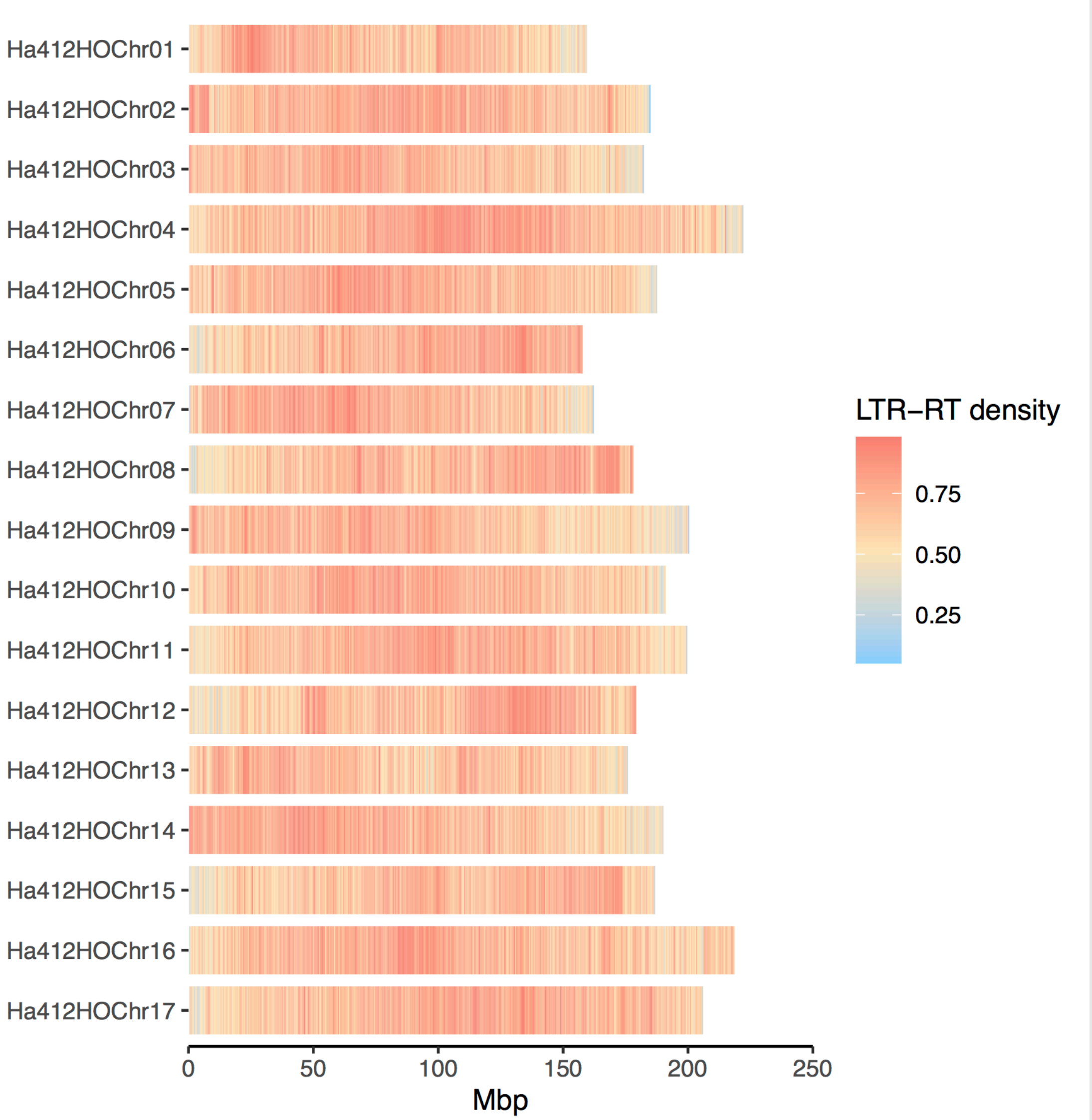


**Figure S7.** Genomic distributions of long terminal repeat retrotransposons (LTR-RTs) in Ha412-HOv2. Density of LTR-RTs was calculated in 500kb bins per chromosome.

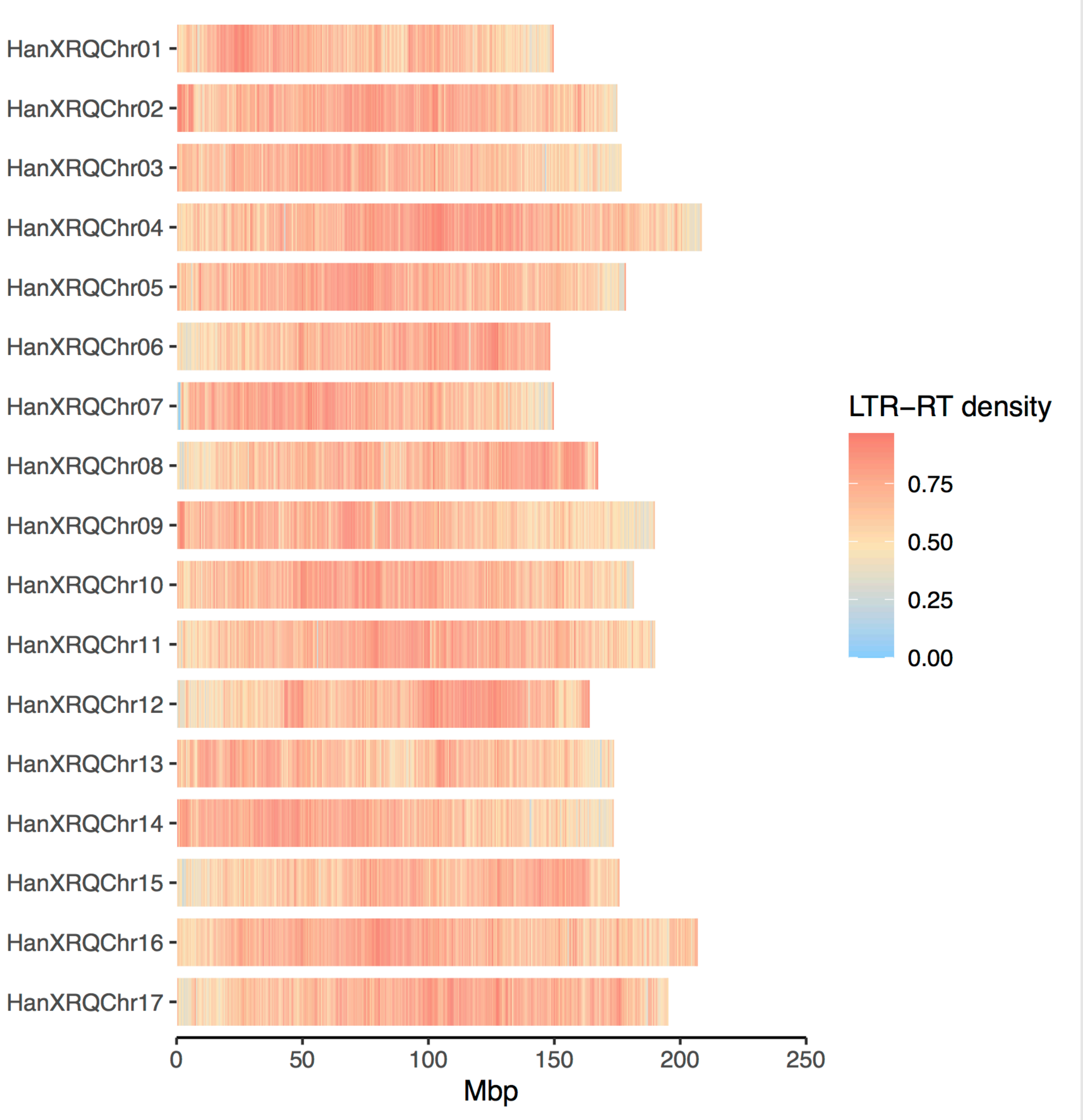


**Figure S8.** Genomic distributions of long terminal repeat retrotransposons (LTR-RTs) in XRQv2. Density of LTR-RTs was calculated in 500kb bins per chromosome.

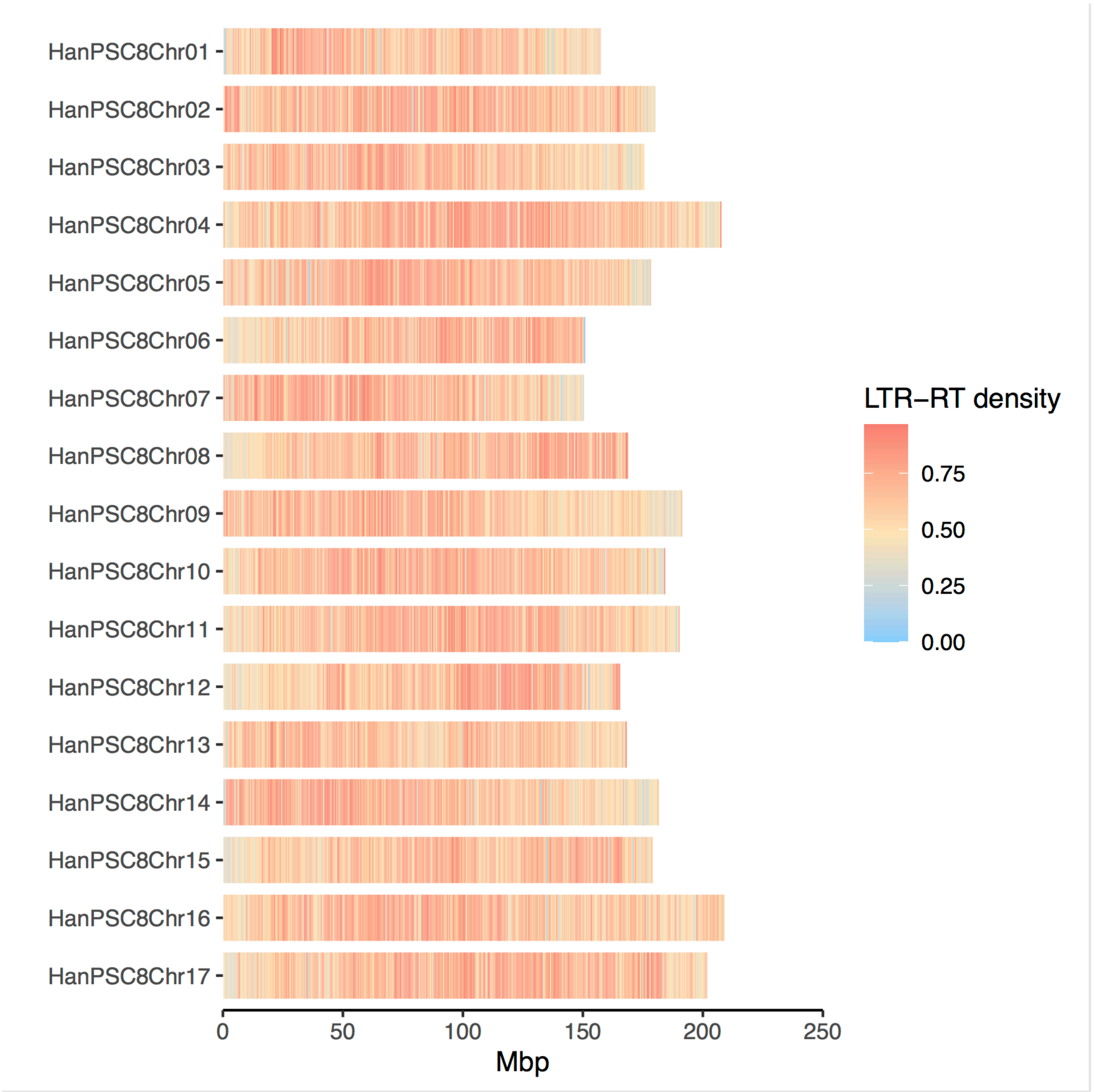


**Figure S9.** Genomic distributions of long terminal repeat retrotransposons (LTR-RTs) in PSC8. Density of LTR-RTs was calculated in 500kb bins per chromosome.

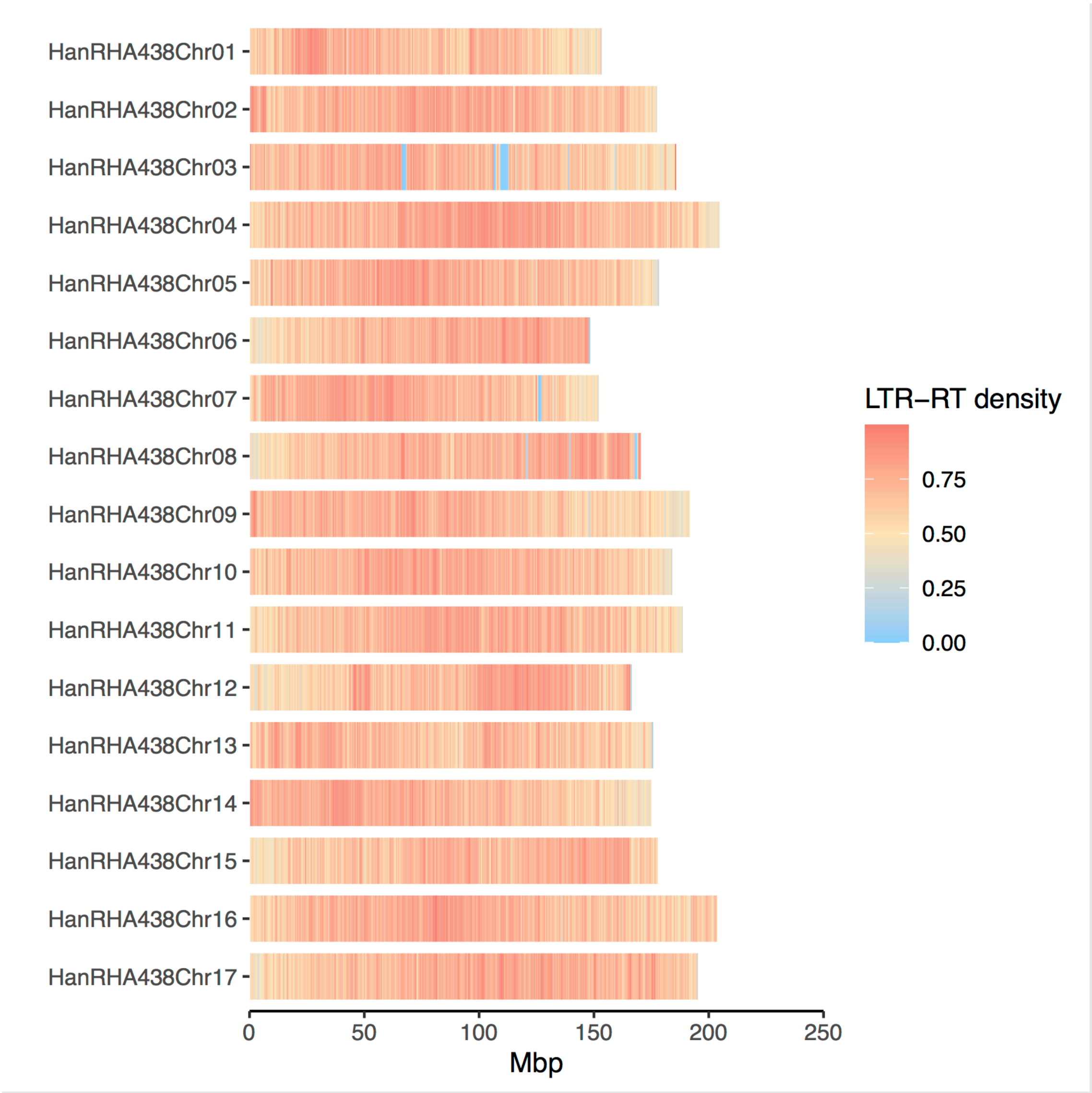


**Figure S10.** Genomic distributions of long terminal repeat retrotransposons (LTR-RTs) in RHA438. Density of LTR-RTs was calculated in 500kb bins per chromosome.

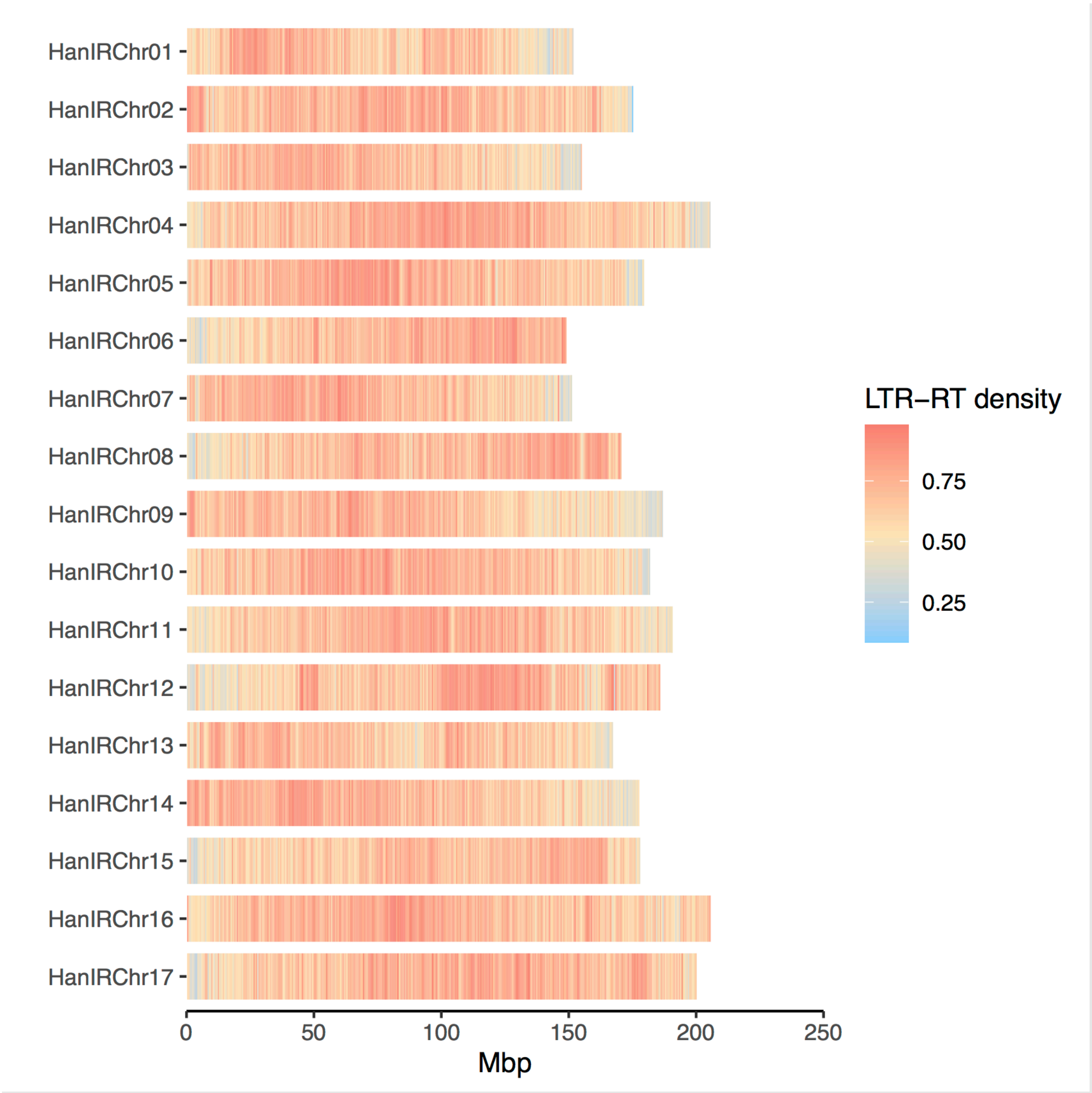


**Figure S11.** Genomic distributions of long terminal repeat retrotransposons (LTR-RTs) in IR. Density of LTR-RTs was calculated in 500kb bins per chromosome.

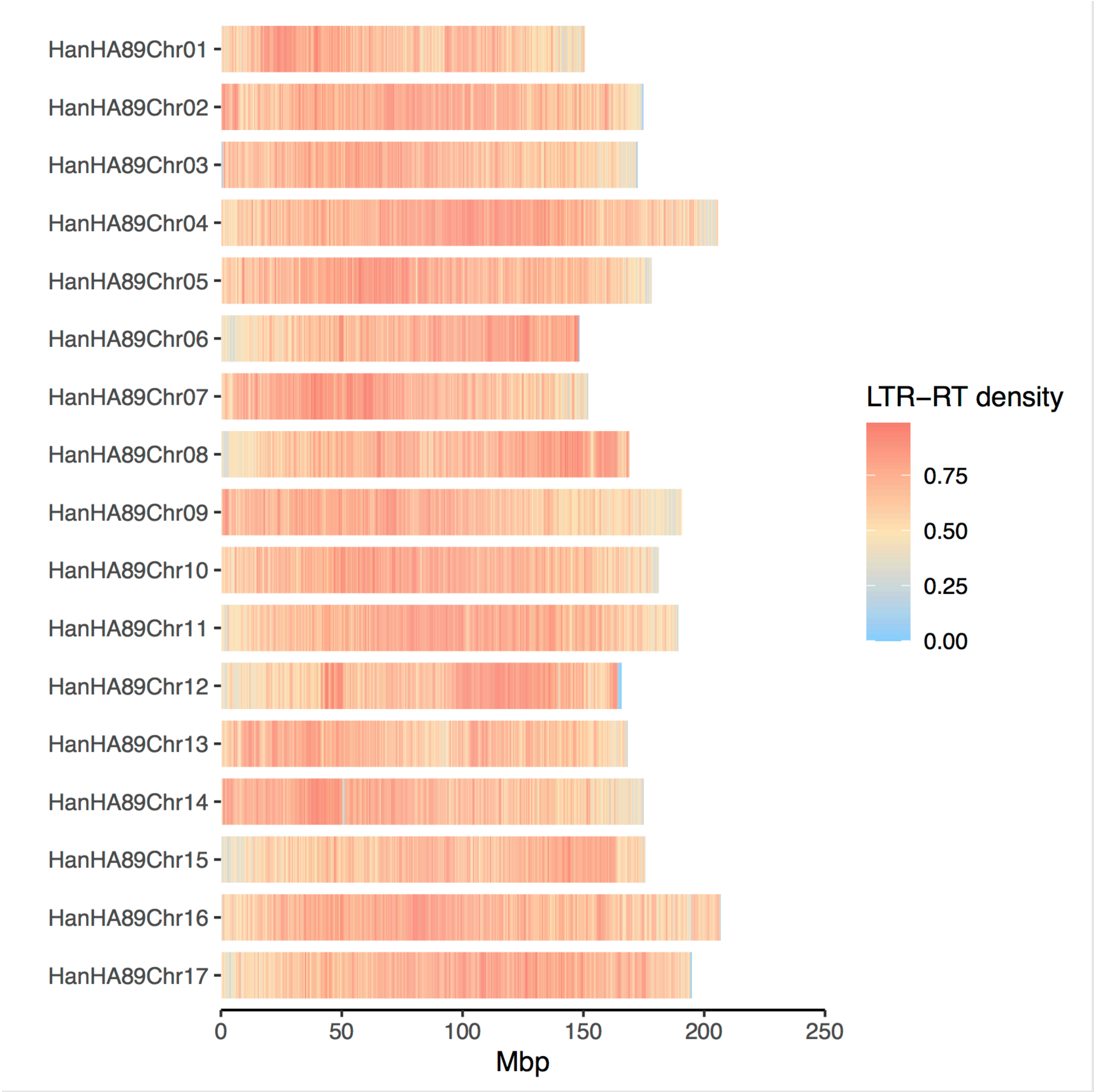


**Figure S12.** Genomic distributions of long terminal repeat retrotransposons (LTR-RTs) in HA89. Density of LTR-RTs was calculated in 500kb bins per chromosome.

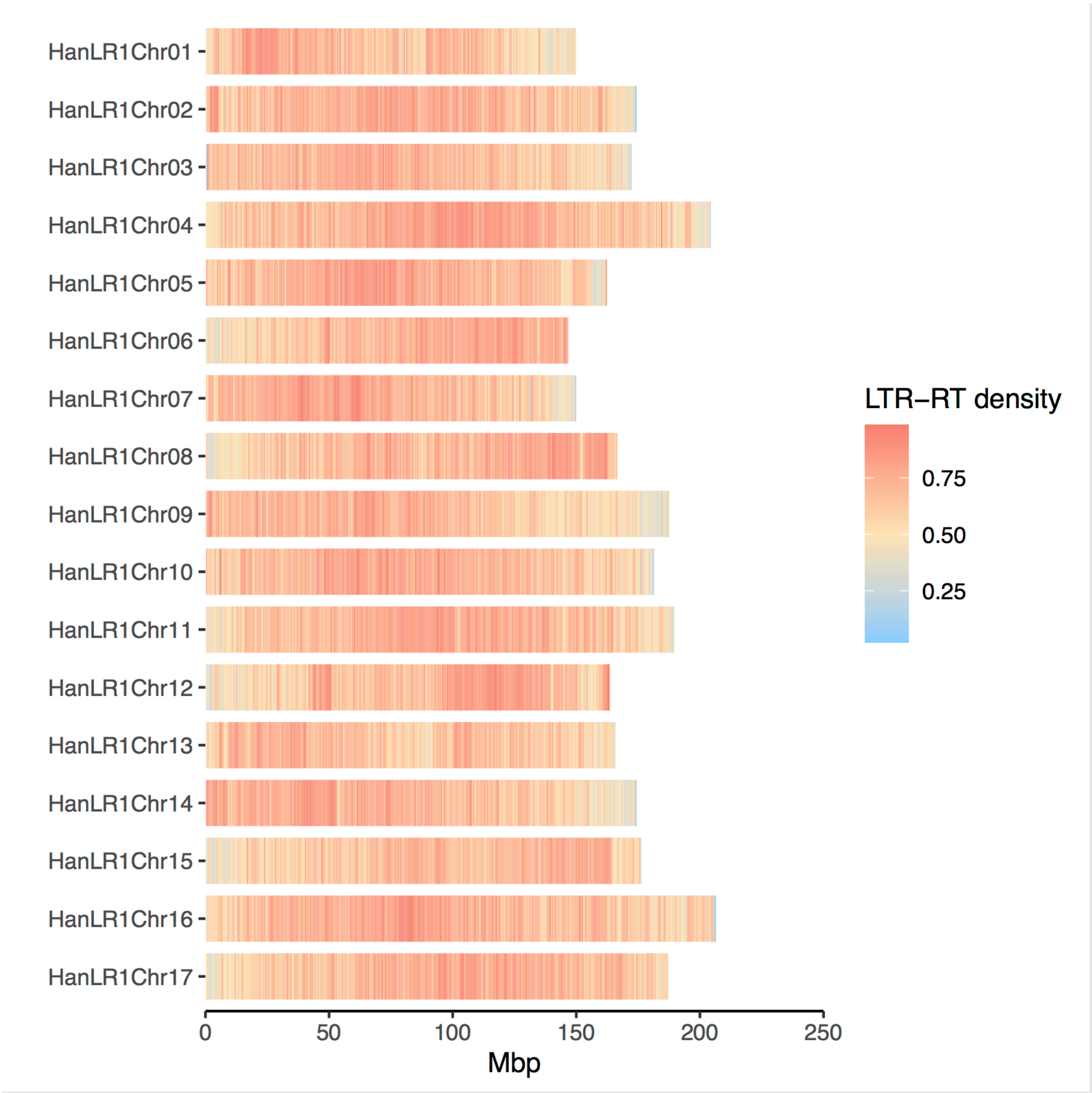


**Figure S13.** Genomic distributions of long terminal repeat retrotransposons (LTR-RTs) in LR1. Density of LTR-RTs was calculated in 500kb bins per chromosome.

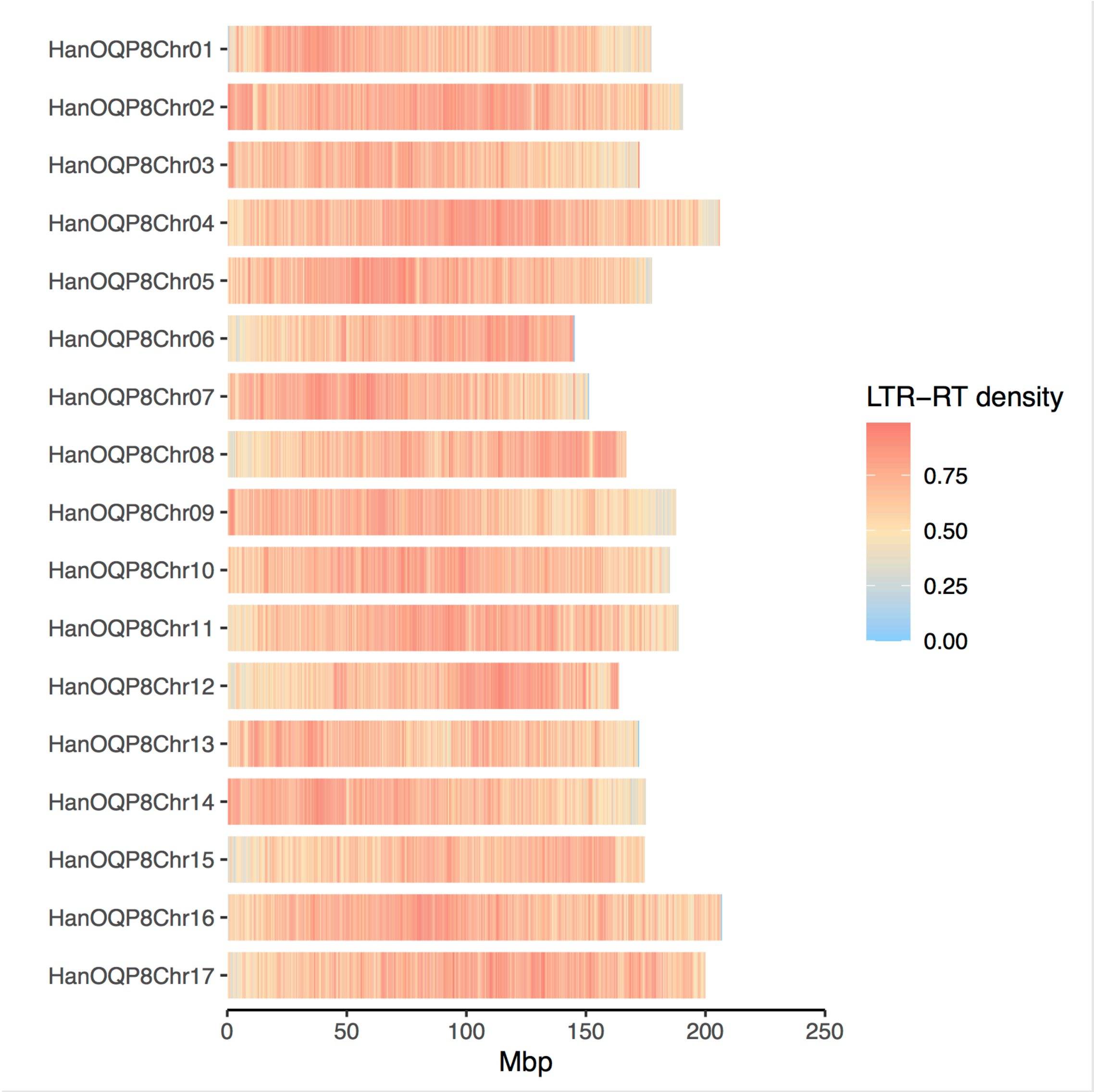


**Figure S14.** Genomic distributions of long terminal repeat retrotransposons (LTR-RTs) in OQP8. Density of LTR-RTs was calculated in 500kb bins per chromosome.

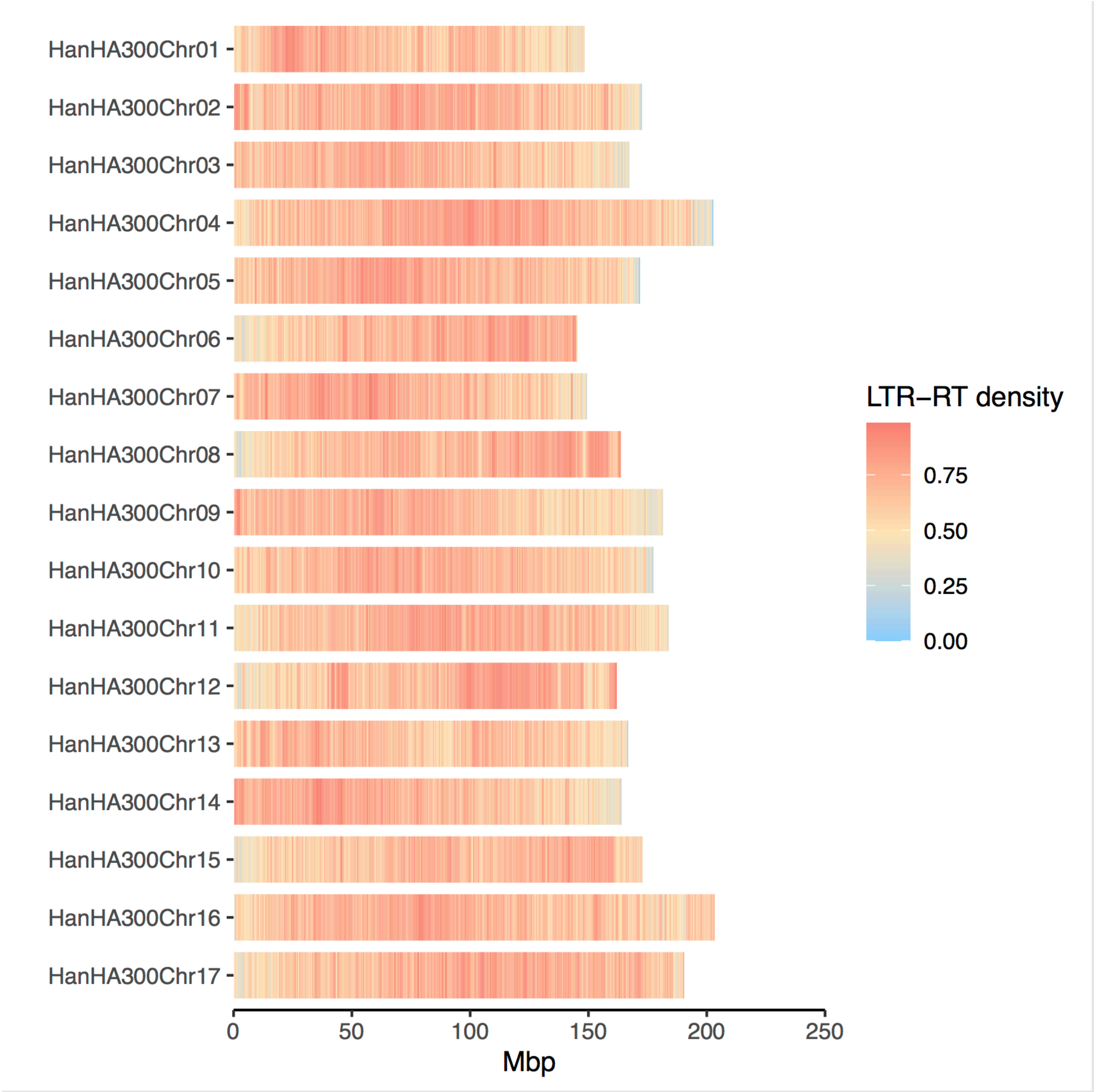


**Figure S15.** Genomic distributions of long terminal repeat retrotransposons (LTR-RTs) in HA300. Density of LTR-RTs was calculated in 500kb bins per chromosome.

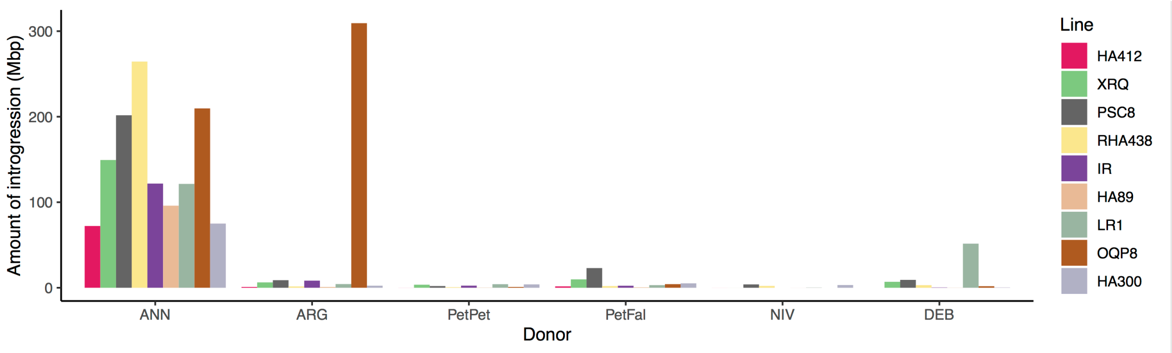


**Figure S16.** Total amount of introgression from each wild donor in the cultivar genomes. ANN: *Helianthus annuus*, ARG: *H. argophyllus*, PetPet: *H. petiolaris* subsp. *petiolaris*, PetFal: *H. petiolaris* subsp. *fallax*, NIV: *H. niveus* and DEB: *H. debilis*.

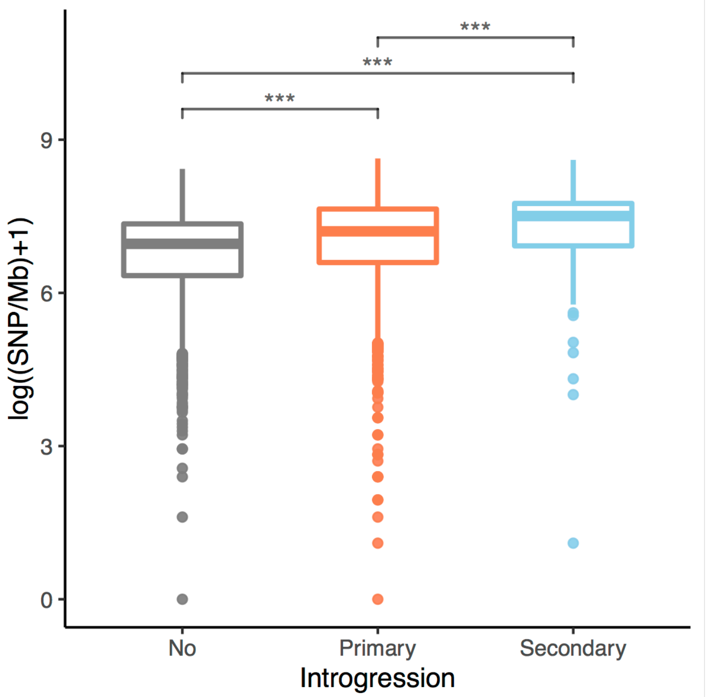


**Figure S17.** Densities of SNPs in regions without introgression, regions with introgressions from the primary gene pool (primary introgressions) and regions from the secondary gene pool (secondary introgression) in the cultivated sunflower association mapping population. The densities of SNPs were calculated in non-overlapping windows of 500kb. Asterisks denote significance in independent t-tests: ***p<0.001.

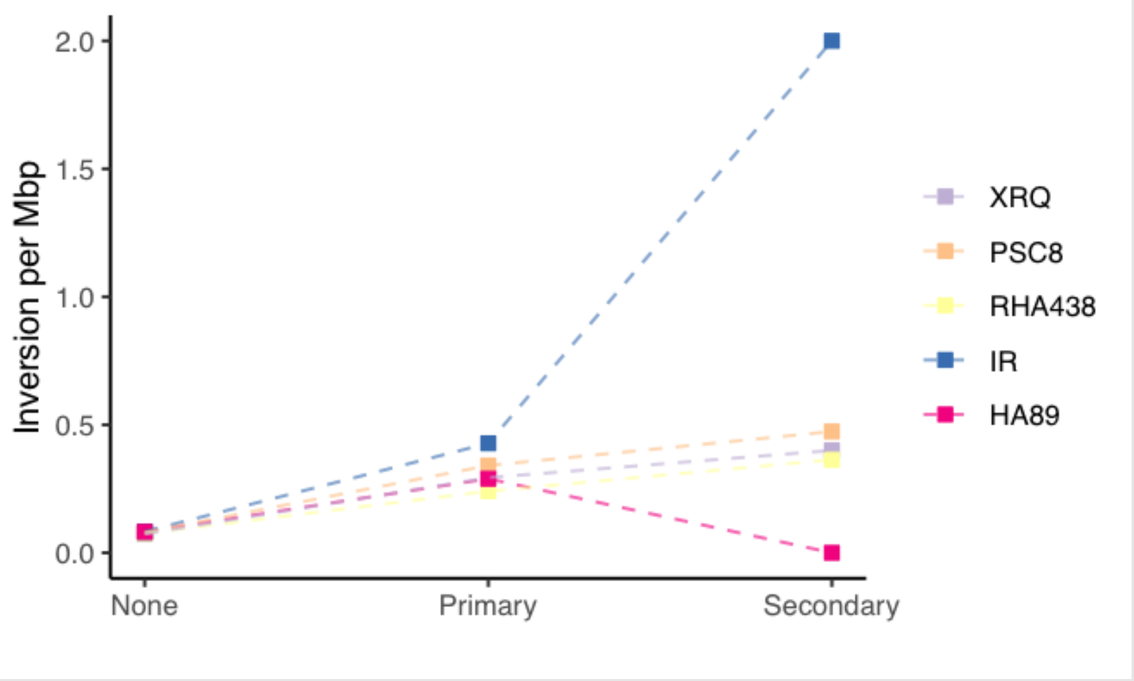


**Figure S18.** Incidence of chromosomal inversions between each of the high-contiguity cultivar genomes and the HA412-HOv2 reference in regions without introgression, regions with introgressions from the primary gene pool (primary introgressions) and regions from the secondary gene pool (secondary introgression).

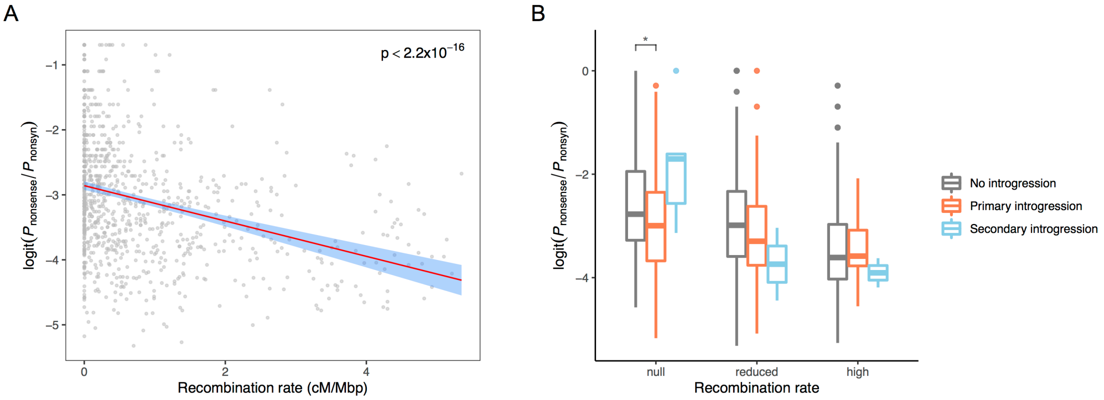


**Figure S19.** Ratio of alternative stop codons and nonsynonymous mutations ($P_{\text{nonsense}}/P_{\text{nonsyn}}$) found in the genome assemblies. **A.** Recombination rate and $P_{\text{nonsense}}/P_{\text{nonsyn}}$. The red lines denote the best-fit linear regression line with the 95% confidence intervals shaded in blue. **B.** $P_{\text{nonsense}}/P_{\text{nonsyn}}$ in regions without introgression, regions with introgressions from the primary gene pool (primary introgressions) and regions from the secondary gene pool (secondary introgression). $P_{\text{nonsense}}/P_{\text{nonsyn}}$ was calculated in non-overlapping windows of 500kb. Windows of each recombination rate category (high: > 2 cM/Mb, reduced: 0.01-2 cM/Mb, null: <0.01 cM/Mb) were compared separately. Asterisks denote significance in independent t-test: *0.05>p>0.01.

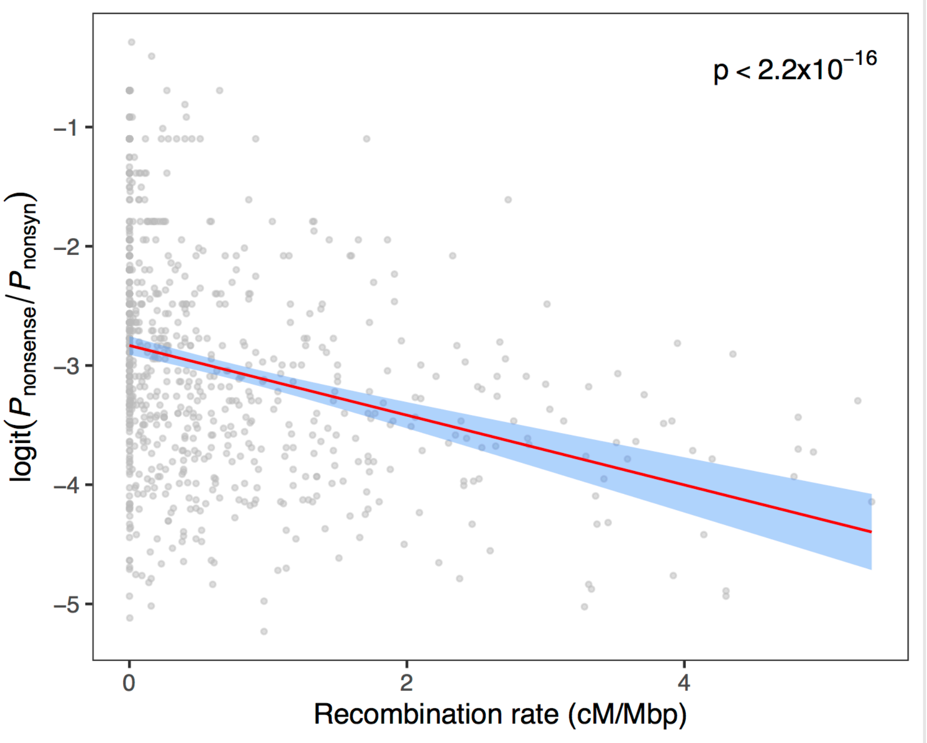


**Figure S20.** Recombination rate and ratio of alternative stop codons and nonsynonymous mutations ($P_{\text{nonsense}}/P_{\text{nonsyn}}$) in the cultivated sunflower association mapping population. The red lines denote the best-fit linear regression line with the 95% confidence intervals shaded in blue.

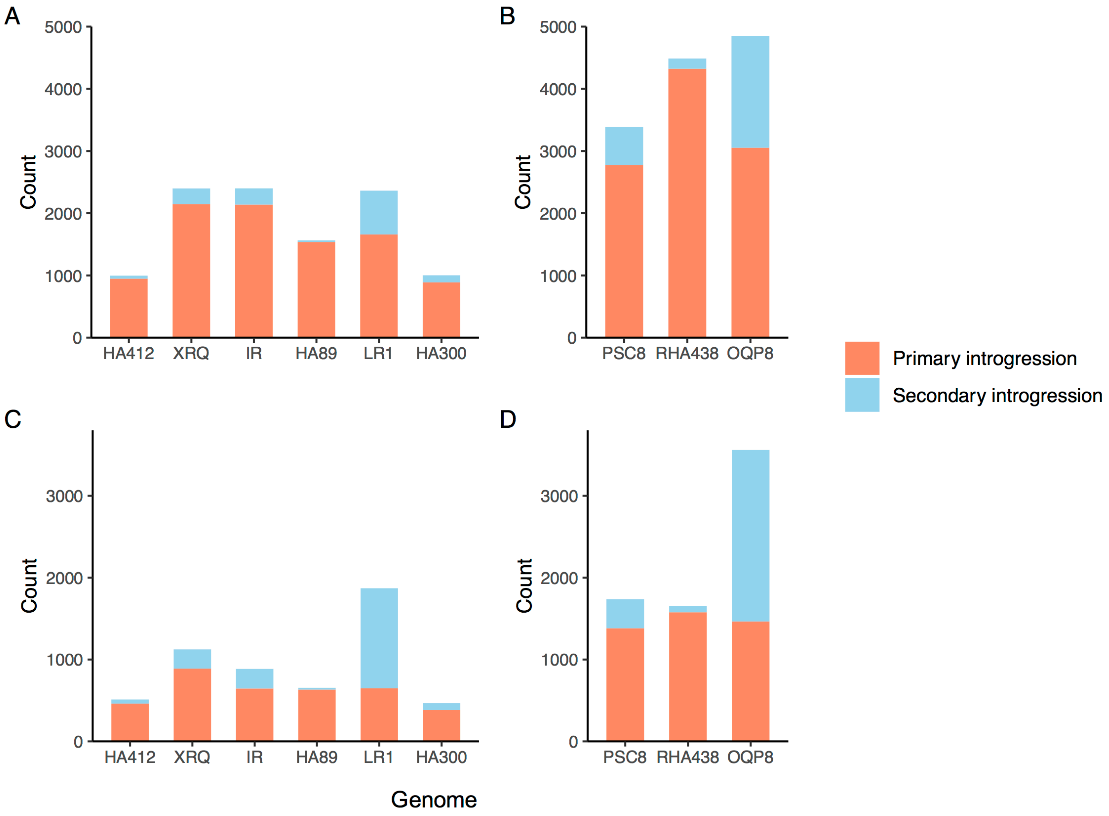


**Figure S21.** Numbers of introduced genes in **A.** maintainer and **B.** restorer lines and numbers of missing genes in **C.** maintainer and **D.** restorer lines.

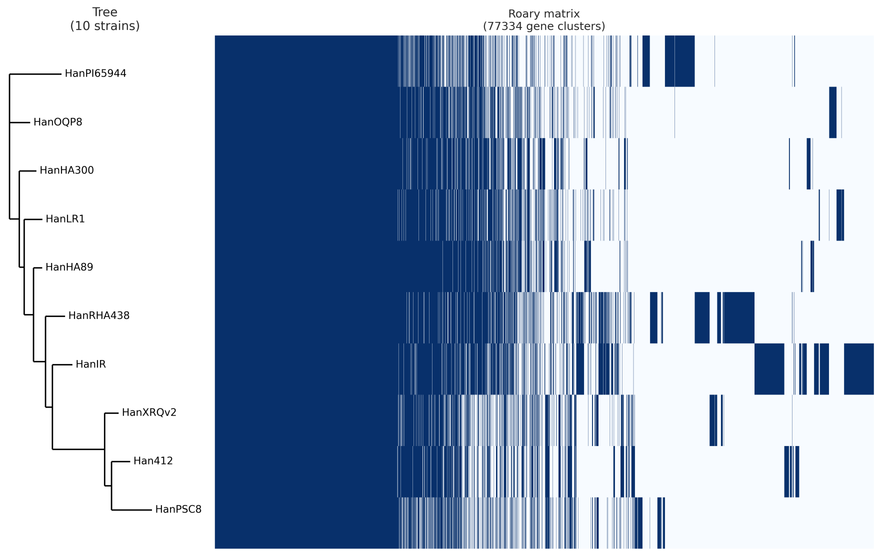


**Figure S22**. Dendrogram of completeness and core genome content in each genome. Core genome content and singletons (unique genes) before GMAP completeness procedure for determination of the final pan-genome PAV table.

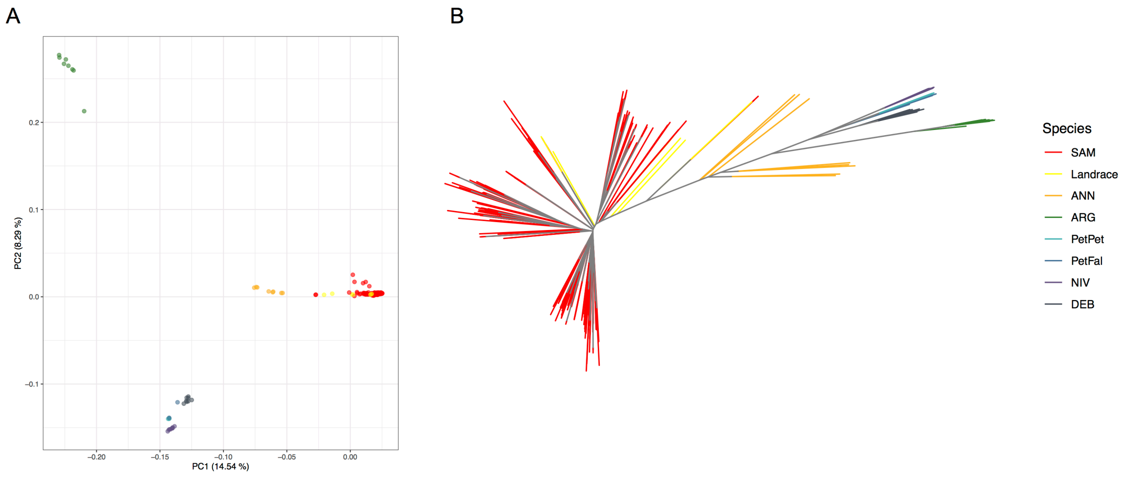


**Figure S23**. **A.** Principal component analysis and **B.** Neighbor-joining tree of 287 samples in the SAM population and 48 landrace and wild samples. ANN: *Helianthus annuus*, ANG: *H. argophyllus*, PetPet: *H. petiolaris* subsp. *petiolaris*, PetFal: *H. petiolaris* subsp. *fallax*, NIV: *H. niveus*, DEB: *H. debilis*.

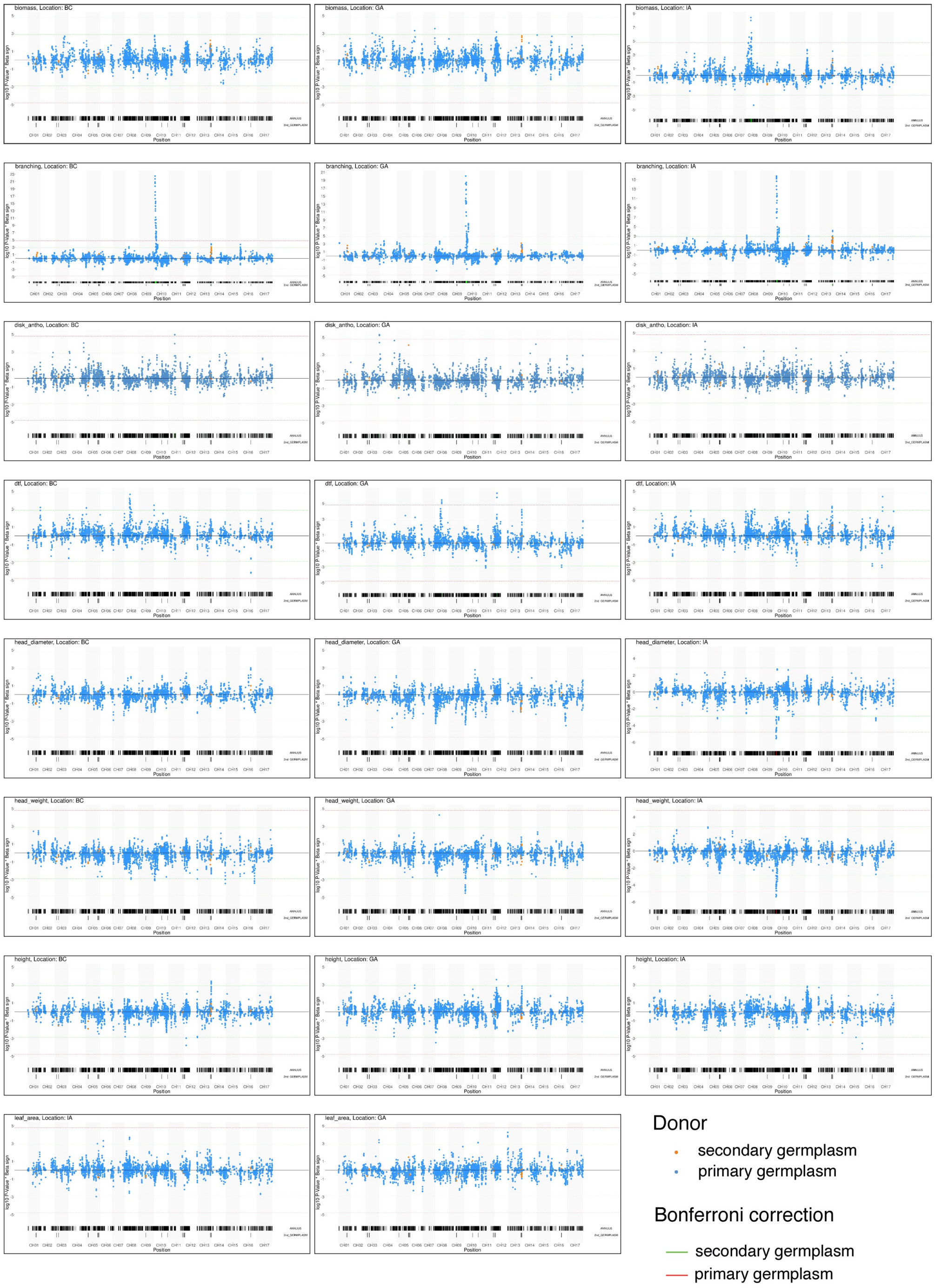
biomass, Location: BC
biomass, Location: GA
biomass, Location: IA
branching, Location: BC
branching, Location: GA
branching, Location: IA
disk_antho, Location: BC
disk_antho, Location: GA
disk_antho, Location: IA
dtf, Location: BC
dtf, Location: GA
dtf, Location: IA
head_diameter, Location: BC
head_diameter, Location: GA
head_diameter, Location: IA
head_weight, Location: BC
head_weight, Location: GA
head_weight, Location: IA
height, Location: BC
height, Location: GA
height, Location: IA
leaf_area, Location: IA
leaf_area, Location: GA
Position
Donor
secondary germplasm
primary germplasm
Bonferroni correction
secondary germplasm
primary germplasm

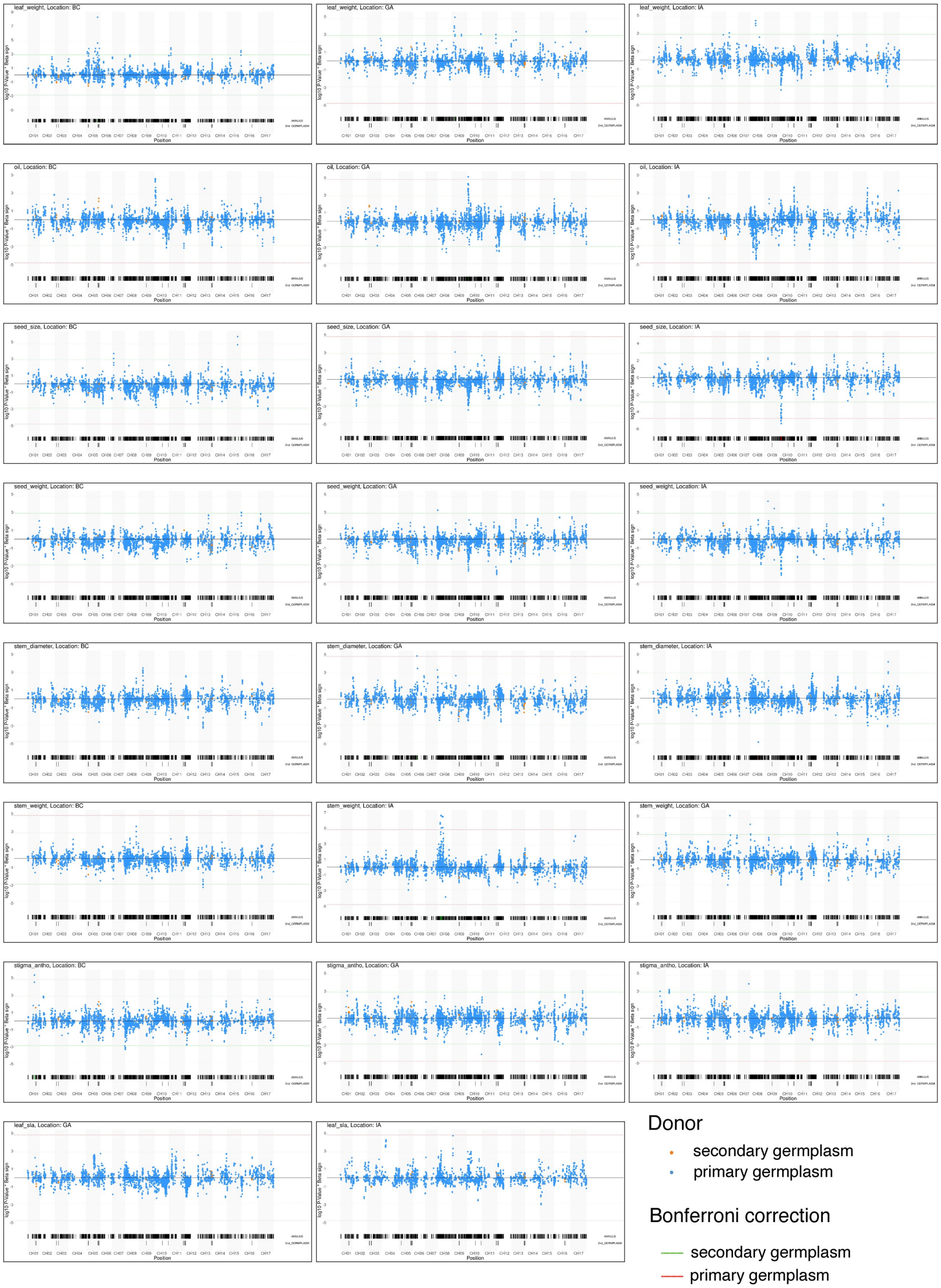


**Figure S24.** Manhattan plots for GWA analyses between introgressions and phenotypic traits in three locations. Blue and orange dots denote introgressions from primary and secondary germplasm, respectively. The red line indicates the Bonferroni-corrected significance threshold for GWAS in the primary introgression dataset, whereas the green line represents the Bonferroni-corrected significance threshold for GWAS in the secondary introgression dataset.

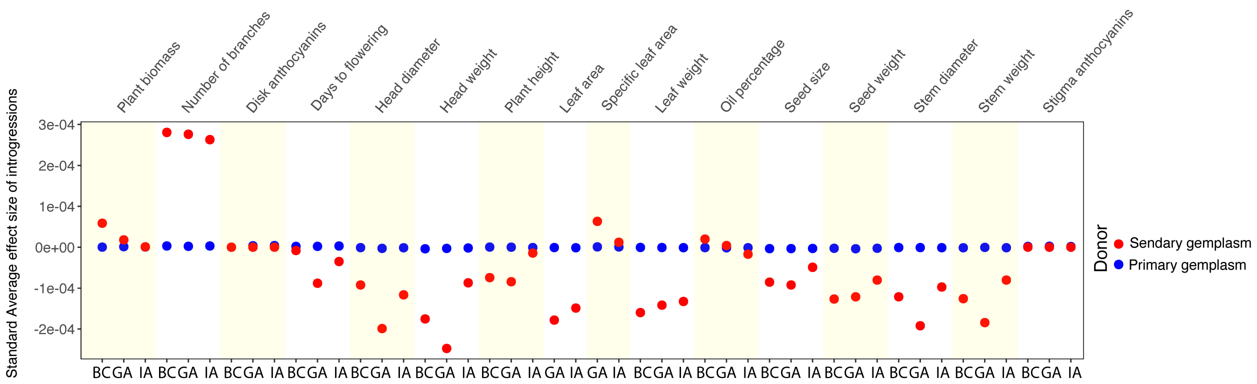


**Figure S25.** The standard average effects of introgression variants (calculated from Z-score normalized trait values) in the SAM population. BC: British Columbia, GA: Georgia, IA: Iowa.

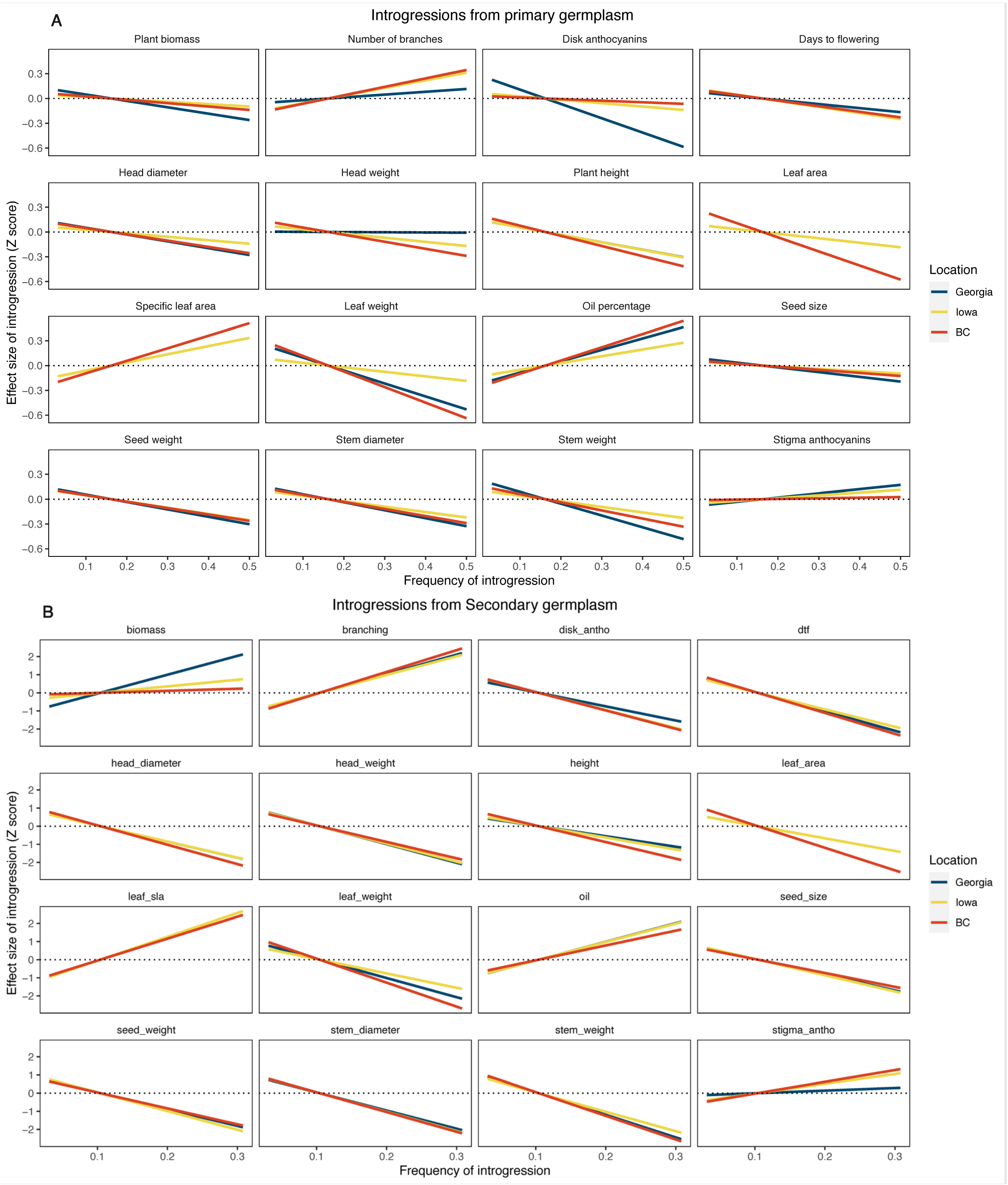


**Figure S26.** A fitted linear regression line between introgression frequency and introgression effect on phenotype variation in the SAM population. **A.** Introgressions from primary germplasm. **B.** Introgressions from secondary germplasm.

**Table S1.** Origin, pedigree, and breeding information about sequenced lines.

| Line & PI code | CMS & branching type | Oil type | Origin, date | Pedigree | Comments from GRIN and INRAE databases |
|---|---|---|---|---|---|
| HA412-HO, PI_642777 | Maintainer, unbranched | Oleic | USDA, 2006[c] | BC4F3 maintainer line selected from HA412*5/HA434 | Derived from backcross between HA412 (Sclerotinia head & stalk rot tolerant) and HA434 (high-oleic maintainer line); Miller et al. (2006). |
| XRQ | Maintainer, unbranched | Linoleic | INRAE, 1996[b] | QPD.(HA89 PROGR)B.1.2.7.C3..2.1 | Bred for downy mildew resistance. |
| PSC8 | Restorer, apical branching | Linoleic | INRAE, 1991[b] | N.ISCMR5.313.6.7.7.5 | Bred for Sclerotinia head rot resistance. |
| RHA438 PI_633748 | Restorer, apical branching | Oleic | USDA, 2004[c] | OR.RHA 438 (RHA340xRHA344) | Bred for mid-oleic or high-oleic fatty acid composition and resistance to metalaxyl-insensitive strains of downy mildew. Miller et al. (2004). |
| IR | Maintainer, unbranched | Linoleic | INRAE, 2004[b] | IMPIRA SC4・194.8.6.C11.1.1.1.3.1 | Bred for Sclerotinia head rot resistance. Fixed from Argentinian population. |
| HA89, PI_599773 | Maintainer, unbranched | Linoleic | USDA, 1971[a] | HA 89 aka CD (INRAE) and VNIIMK 8931 | Bred for high oil content and rust resistance. |
| LR1 | Maintainer, unbranched | Linoleic | INRAE, 1989[b] | DEBILIS.215 x 85B6 89-1471-12 | Introgression line from *Helianthus debilis* carrying Orobanche resistance locus. |
| OQP8 | Restorer, apical branching | Oleic | INRAE, 2005[b] | OR.(PAA1 RHA345)A.1.9.18.1.2 | Bred for oleic fatty acids with introgression of downy mildew resistance QTL from *Helianthus argophyllus*. |
| HA300 PI_552938 | Maintainer, unbranched | Linoleic | USDA, 1979[c] | F5 selection from RHA340 (PI 5187778)/RHA344 (PI 509054). | Fertility restorer germplasm advanced by pedigree selection from cross between RHA340 and RHA344. Fick et al. (1979). |
| PI659440 | Wild, branched | N/A | Nebraska, USA | Collection from natural population in Keith Co., NE | Self-compatible wild germplasm inbred for several generations. |

N/A = not available.
[a]Year of registration in GRIN.
[b]Year of introduction to INRAE genebank.
[c]Year of publication.

**Table S2.** Quantity and fragment length of high molecular weight DNA for HA89, LR1, OQP8, and PI659440 genotypes.

| Genotype | Volume (µl) | [ ng/µl] Nanodrop | [ ng/µl] QuBiT | Quantity (µg) DNA | % fragments >20kb |
|---|---|---|---|---|---|
| | 250 | 114 | 152 | ~33 | N/A |
| HA89 | 450 | 133 | 186 | ~71 | 84 |
| | 450 | 153 | 311 | ~104 | 84 |
| LR1 | 200 | 170 | 207 | ~37 | 65 |
| OQP8 | 200 | 188 | 252 | ~44 | 48 |
| HA300 | 150 | 180 | 212 | ~29 | N/A |
| PI659440 | 1000 | 130 | 108 | ~119 | 36 |

N/A = not available.

**Table S3.** Quantity and quality of RNA for the HA89 and PI659440 genotypes.

| Genotype | Volume (µl) | [ng/µl] Nanodrop | RNA Integrity Number (RIN) | Quantity (µg) RNA |
|---|---|---|---|---|
| HA89 - leaf | 40 | 280.6 | 7.2 | ~ 11 |
| HA89 - bud | 40 | 84.4 | 8.8 | ~ 3.3 |
| HA89 - stem | 40 | 120.3 | 8.7 | ~ 4.8 |
| PI659440 - leaf | 40 | 177.4 | 6.5 | ~ 7 |
| PI659440 - bud | 40 | 160.4 | 8.8 | ~ 6.4 |
| PI659440 - stem | 40 | 60.8 | 8.6 | ~ 2.4 |

**Table S4.** Gene finding protocol and gene modeling assessments.

| Genotype | EuGene release | EuGene pipeline release | Transcriptome datasets used in the pipeline (minimum percentage identity >= 99% for IsoSeq datasets) | Complete BUSCOs (%) | Complete & single copy BUSCOs (%) | Complete & duplicated BUSCOs (%) | Fragmented BUSCOs (%) | Missing BUSCOs (%) | Total BUSCO groups searched |
|---|---|---|---|---|---|---|---|---|---|
| HA412-HOv2 | 4.2a | 1.5 | Badouin et al. 2017 + PRJNA517222 IsoSeq | 97.9 | 87.2 | 10.7 | 1.3 | 0.8 | 1614 |
| XRQv2 | 4.2a | 1.5 | Badouin et al. 2017 + PRJNA517222 IsoSeq | 97.4 | 86.7 | 10.7 | 1.3 | 1.3 | 1614 |
| PSC8 | 4.2a | 1.5 | Badouin et al. 2017 + PRJNA517222 IsoSeq | 94.5 | 84.6 | 9.9 | 1.5 | 4 | 1614 |
| RHA438 | 4.2a | 1.6 | RHA438 Isoseq, this work | 96.7 | 85.1 | 11.6 | 1.5 | 1.8 | 1614 |
| IR | 4.2a | 1.6 | IR Isoseq, this work | 97.1 | 85.7 | 11.4 | 1.3 | 1.6 | 1614 |
| HA89 | 4.2a | 1.6-dev | HA89 Isoseq, this work | 97.3 | 87.1 | 10.2 | 1.7 | 1 | 1614 |
| LR1 | 4.2b | 1.6-dev | HA89 Isoseq, this work | 85.9 | 76.3 | 9.6 | 6.3 | 7.8 | 1614 |
| OPQ8 | 4.2b | 1.6-dev | HA89 Isoseq, this work | 88.1 | 79.3 | 8.8 | 8.2 | 3.7 | 1614 |
| HA300 | 4.2b | 1.6-dev | HA89 Isoseq, this work | 90.3 | 81.1 | 9.2 | 6.3 | 3.4 | 1614 |
| PI659440 | 4.2b | 1.6-dev | PI659440 IsoSeq, this work | 96.5 | 83.9 | 12.6 | 2 | 1.5 | 1614 |

**Table S5.** Gene summary data after filtering.

| Assembly | Data | gene | mRNA | exon | 5' UTR | CDS | 3' UTR | Genes dropped |
|---|---|---|---|---|---|---|---|---|
| HA412-HOv2 | RAW data | 87,728 | 72,995 | 308,368 | 63,420 | 255,787 | 61,951 | -40,960 |
| | Post_Filtering | 46,768 | 45,008 | 234,380 | 47,293 | 202,009 | 45,624 | |
| XRQv2 | RAW data | 83,726 | 71,289 | 304,285 | 63,403 | 252,381 | 62,882 | -37,682 |
| | Post_Filtering | 46,044 | 44,640 | 233,219 | 47,479 | 200,724 | 46,407 | |
| PSC8 | RAW data | 80,979 | 68,610 | 291,253 | 60,526 | 241,974 | 58,430 | -36,815 |
| | Post_Filtering | 44,164 | 42,766 | 221,752 | 45,136 | 191,508 | 42,908 | |
| RHA438 | RAW data | 86,603 | 75,639 | 307,086 | 77,413 | 257,336 | 73,963 | -25,693 |
| | Post_Filtering | 60,910 | 59,592 | 272,088 | 65,835 | 230,301 | 63,054 | |
| IR | RAW data | 91,844 | 80,741 | 319,322 | 84,915 | 263,682 | 81,317 | -29,101 |
| | Post_Filtering | 62,743 | 61,109 | 277,213 | 69,983 | 231,833 | 66,959 | |
| HA89 | RAW data | 75,799 | 62,812 | 287,277 | 61,517 | 239,245 | 57,554 | -22,529 |
| | Post_Filtering | 53,270 | 50,203 | 257,058 | 54,203 | 215,942 | 50,900 | |
| LR1 | RAW data | 88,001 | 65,594 | 271,989 | 46,309 | 236,889 | 43,341 | -25,961 |
| | Post_Filtering | 62,040 | 51,899 | 242,285 | 40,827 | 212,328 | 38,304 | |
| OQP8 | RAW data | 86,175 | 60,613 | 244,739 | 34,124 | 220,020 | 32,482 | -38,366 |
| | Post_Filtering | 47,809 | 47,287 | 217,269 | 30,520 | 196,105 | 29,160 | |
| HA300 | RAW data | 84,348 | 63,048 | 266,478 | 47,398 | 230,417 | 44,772 | -34,845 |
| | Post_Filtering | 49,503 | 48,777 | 235,743 | 41,361 | 205,259 | 39,265 | |
| PI659440 | RAW data | 75,639 | 61,323 | 264,710 | 49,004 | 234,047 | 44,374 | -25,528 |
| | Post_Filtering | 50,111 | 48,448 | 237,001 | 43,028 | 211,013 | 39,310 | |

**Table S6**. Summary of transposable elements in each genome. Size (bp) and proportion of each class of transposable elements were summarized across 17 chromosomes.

| **Assembly** | **LTR** | | | **TIR** | | | | | **non-TIR** | **Total** |
|---|---|---|---|---|---|---|---|---|---|---|
| | ***Copia*** | ***Gypsy*** | **unknown** | ***CACTA*** | ***Mutator*** | ***PIF-Harbinger*** | ***Tc1-Mariner*** | ***hAT*** | ***Helitron*** | |
| HA412-HOv2 | 350,802,612 | 1,326,002,146 | 578,541,122 | 26,488,425 | 113,888,703 | 14,911,284 | 1,570,160 | 90,188,880 | 115,857,160 | 2,618,250,492 |
| | 11.12% | 42.04% | 18.34% | 0.84% | 3.61% | 0.47% | 0.05% | 2.86% | 3.67% | 83.02% |
| XRQv2 | 353,994,515 | 1,232,379,833 | 506,296,443 | 28,798,944 | 118,879,762 | 15,602,396 | 1,726,118 | 80,285,030 | 111,438,725 | 2,449,401,766 |
| | 12.04% | 41.93% | 17.23% | 0.98% | 4.05% | 0.53% | 0.06% | 2.73% | 3.79% | 83.33% |
| PSC8 | 346,234,534 | 1,151,044,163 | 474,037,039 | 26,490,261 | 105,156,622 | 14,783,567 | 1,885,190 | 74,165,631 | 107,245,364 | 2,301,042,371 |
| | 12.52% | 41.63% | 17.14% | 0.96% | 3.80% | 0.53% | 0.07% | 2.68% | 3.88% | 83.21% |
| RHA438 | 407,699,019 | 1,347,612,222 | 444,214,721 | 3,428,148 | 13,477,334 | 920,658 | 230,667 | 8,313,021 | 12,254,387 | 2,238,150,177 |
| | 13.56% | 44.83% | 14.78% | 0.11% | 0.45% | 0.03% | 0.01% | 0.28% | 0.41% | 74.45% |
| IR | 375,988,121 | 1,302,978,999 | 424,700,560 | 16,634,950 | 74,135,512 | 9,484,702 | 1,787,266 | 52,259,062 | 78,686,600 | 2,336,655,772 |
| | 12.52% | 43.37% | 14.14% | 0.55% | 2.47% | 0.32% | 0.06% | 1.74% | 2.62% | 77.78% |
| HA89 | 377,815,177 | 1,307,727,359 | 412,441,433 | 16,181,369 | 71,775,461 | 8,840,313 | 1,530,347 | 48,780,825 | 84,070,056 | 2,329,162,340 |
| | 12.62% | 43.69% | 13.78% | 0.54% | 2.40% | 0.30% | 0.05% | 1.63% | 2.81% | 77.82% |
| LR1 | 371,329,621 | 1,293,395,471 | 409,482,041 | 16,324,640 | 73,797,667 | 8,995,062 | 1,418,773 | 51,429,438 | 73,241,890 | 2,299,414,603 |
| | 12.57% | 43.77% | 13.86% | 0.55% | 2.50% | 0.30% | 0.05% | 1.74% | 2.48% | 77.82% |
| OQP8 | 375,899,400 | 1,345,429,323 | 420,593,182 | 15,875,969 | 74,521,490 | 9,873,865 | 1,290,921 | 51,606,736 | 81,374,064 | 2,376,464,950 |
| | 12.37% | 44.29% | 13.85% | 0.52% | 2.45% | 0.33% | 0.04% | 1.70% | 2.68% | 78.23% |
| HA300 | 365,625,093 | 1,275,401,277 | 405,596,471 | 15,201,266 | 74,229,615 | 8,974,337 | 1,452,000 | 50,290,939 | 79,000,215 | 2,275,771,213 |
| | 12.52% | 43.69% | 13.89% | 0.52% | 2.54% | 0.31% | 0.05% | 1.72% | 2.71% | 77.96% |
| PI659440 | 377,184,453 | 1,313,298,182 | 422,637,602 | 18,049,679 | 77,921,012 | 8,798,722 | 1,900,566 | 49,443,044 | 81,569,963 | 2,350,803,223 |
| | 12.50% | 43.53% | 14.01% | 0.60% | 2.58% | 0.29% | 0.06% | 1.64% | 2.70% | 77.92% |

**Table S7.** Introgressed genomic intervals associated with traits in GWA analysis. Interval coordinates are based on the HA412-HOv2 reference.

| Donor | Introgression effect | Trait | Location | | |
|---|---|---|---|---|---|
| | | | BC | GA | IA |
| Primary | increase | Biomass | - | - | Chr08:42030001-42255000, Chr08:64812001-72109000 |
| Primary | increase | Branching | Chr10:12812001-29489000 | Chr10:12812001-29488000, Chr10:46392001-47108000 | Chr10:12812001-24575000 |
| Primary | increase | Disk anthocyanins | Chr11:80778001-82491000 | Chr04:3604001-3713000 | - |
| Primary | increase | Days to flower | - | Chr08:84431001-84484000, Chr12:29515001-29569000 | - |
| Primary | increase | Leaf weight | Chr05:151464001-151465000 | Chr09:44584001-44585000 | - |
| Primary | increase | Oil content | - | Chr10:14741001-15267000 | - |
| Primary | increase | Seed size | Chr15:140714001-140837000 | - | Chr10:13317001-20477000 |
| Primary | increase | Stem diameter | - | Chr06:48820001-49008000 | - |
| Primary | increase | Stem weight | - | Chr06:48820001-49008000 | Chr08:32042001-46640000, Chr08:64812001-65162000 |
| Primary | increase | Stigma anthocyanins | - | Chr01:88822001-89550000 | - |
| Primary | decrease | Head diameter | - | - | Chr10:13317001-17905000 |
| Primary | decrease | Head weight | - | - | Chr10:13139001-17942000 |
| Primary | decrease | Stem diameter | - | - | Chr08:100411001-100514000 |
| Secondary | increase | Branching | Chr13:173707001-173708000 | Chr13:163451001-163452000 | Chr13:163451001-163452000 |
| Secondary | increase | Disk anthocyanins | - | Chr05:159278001-159449000 | - |

Primary = Primary germplasm
Secondary = Secondary germplasm

**Table S8**. NCBI accessions for raw sequence data.

| Genotype | Sequencing technology | NCBI accession |
|---|---|---|
| XRQ | ILLUMINA | PRJNA345532 |
| PSC8 | RSII | SRP373016 |
| PSC8 | ILLUMINA | SRP373015 |
| RHA438 | Genomic Sequel | SRP373014 |
| RHA438 | IsoSeq | SRP373269 |
| HA89 | IsoSeq | SRP373019 |
| HA89 | Genomic HiFi | SRP373018 |
| LR1 | Genomic HiFi | SRP373020 |
| HA300 | Genomic HiFi | SRP373017 |
| IR | Genomic HiFi | SRP373013 |
| IR | IsoSeq | SRP373521 |
| OQP8 | Genomic HiFi | SRP373021 |
| PI659440 | IsoSeq | SRP373523 |
| PI659440 | Genomic HiFi | SRP373522 |
